# Subfunctionalization of *tbx2* paralogues during photoreceptor cell specification in zebrafish

**DOI:** 10.64898/2026.07.01.735836

**Authors:** Austin M. Werner, Jacob A. Dilliplane, Karen Alvarez-Delfin, Michèle G. DuVal, W. Ted Allison, Fan Xiu Zhu, James M. Fadool

## Abstract

Zebrafish possess three distinct sources of retinal progenitors that produce identical photoreceptor subtypes throughout life. All photoreceptor progenitors simultaneously express multiple transcription factors specifying different identities, requiring mechanisms to repress alternative fates. Disruption of the *tbx2* paralogues, *tbx2a* or *tbx2b*, resulted in a cell-fate switch of sws1 cones into rods. Here, we demonstrate that *tbx2b* was necessary for sws1 cone differentiation during embryogenesis and outgrowth at the retinal margin, but *tbx2a* was necessary during photoreceptor regeneration. Transgenic overexpression of Tbx2b was not sufficient to drive the sws1 cone fate or sws1 opsin expression. Rather, Tbx2b repressed the synergistic activity of Nrl and Crx at the rhodopsin promoter. Targeting the transcription factor *thrβ2* on wildtype and *tbx2* mutant backgrounds revealed a hierarchy wherein early progenitors have the potential to be respecified from lws cones into sws1 cones or rods. But late progenitors are limited to either the sws1 cone or rod fate. These data support a model in which transcriptional repressors, like *tbx2a* and *tbx2b,* orchestrate progression through competency states.

## Introduction

Zebrafish provide a tractable model to investigate neurogenesis across lifespan. During embryogenesis, eye field specification, neurogenesis, and circuit formation are highly conserved across species. To match continuous eye growth throughout the life of teleost, new neurons, including photoreceptors, are added at the retinal periphery from a population of mitotic progenitors located in a stem cell niche within the ciliary marginal zone (CMZ). Additionally, rod photoreceptors are continually generated across the central retinal by mitotic rod progenitors, which are derived from a population of slowly dividing neurogenic radial glial cells, the Muller glia (Johns and Fernald, 1981; Raymond et al. 2006; Lenkowski and Raymond, 2014). Lastly, following acute or chronic damage, Muller glia derived mitotic progenitors have the capacity to regenerate all missing cell types, including photoreceptors, and restore vision (Cameron and Easter, 1995; Vihtelic and Hyde, 2000; McGinn et al. 2018).

Photoreceptor specification is tightly regulated by the precise spatial and temporal expression of transcription factors. Most extant vertebrates possess two major classes of retinal photoreceptors, cones and rods. Cones function in bright light, mediate color vision and are subdivided by opsin expression and morphology. Rods express the visual pigment rhodopsin and function in dim light conditions Many teleost, reptiles, and birds possess rods and 4 distinct cone subtypes with peak sensitivity to ultraviolet or violet (SWS1), blue (SWS2), green (RH2), or red (LWS) wavelengths of light. Most mammals possess a rod-dominated retina and retain only two of the four ancestral cone subtypes. The initial steps of photoreceptor specification are highly conserved across vertebrates. Rod and cone precursors share expression of the transcription factors *Orthodenticle homeobox 2 (Otx2)* and *Cone rod homeobox (Crx)* (Furukuwa et al. 1997; Nishida et al. 2003). Current models portray cell-specific transcription factors as binary switches, driving bipotent progenitors into one of two alternative photoreceptor fates. *Thyroid hormone receptor beta 2 (Thrβ2)* is necessary and sufficient for specification of the LWS cone fate (Ng et al. 2001; Ng et al. 2011; Roberts et al. 2006; Glaschke et al. 2011; Suzuki et al. 2013; Deveau et al. 2020; Volkov et al. 2020), whereas expression of *Neural retina leucine zipper* (*Nrl*) specifies the rod fate (Mears et al. 2001; Oh et al. 2007; Oh et al. 2008; Oel et al. 2020). In the absence of *Thrβ2* or *Nrl*, progenitors differentiate as SWS1 cones, leading to the hypothesis that SWS1 cones are the default photoreceptor subtype (Ng et al. 2011; Aramaki et al. 2022). Small changes in the expression of developmentally regulated genes are a potential source of the variation in the number, type, and arrangement of photoreceptors across species (Oh et al. 2008; Sotolongo-Lopez et al. 2016; Volkov et al. 2024; Weit et al. 2026).

Challenging the notion of a default photoreceptor cell fate, we identified in zebrafish that mutations of *t-box transcription factor 2b (tbx2b)* result in a cell fate switch of sws1 cones into rods (Alvarez-Delfin et al. 2009). The ancestors to all extant teleosts underwent a whole genome duplication, and data suggest that approximately thirty percent of the duplicate genes and chromosome segments were retained (Postlewait et al. 2000), including two ohnologues of *tbx2*, *tbx2a and tbx2b*. Gene expression analysis showed that both were expressed in several tissues and multiple cell types in the retina, including the retinal progenitors, ganglion cells, and mature sws1 and sws2 cones (Farrell et al. 2018; Ogawa et al. 2021; Ogawa and Corbo, 2021; Angueyra et al. 2023; Sur et al. 2023). Knockdown of *tbx2a* in embryonic zebrafish resulted in a modest reduction in sws1 cones and a slight increase in rods, suggesting that *tbx2a* and *tbx2b* have overlapping functions (Angueyra et al. 2023). Moreover, knockdown of *tbx2a* or *tbx2b* resulted in dysregulation of Rh2 opsin expression in lws and sws2 cones, respectively (Angueyra et al. 2023). In the developing chick retina, *tbx2* is expressed in sws1 cones (Enright et al. 2015), and in mice, overexpression of *Tbx3* led to a reduction in rods supporting conserved roles of t-box transcription factors across species. Most evidence supports Tbx2 acts as a transcriptional suppressor in development and cancer posing new questions regarding its roles in photoreceptor development (Zhu et al. 2014; Cho et al. 2017; Teegala et al. 2018).

Using gene targeting approaches, we uncovered shifting roles for *tbx2a* and *tbx2b* in different populations of retinal progenitors. Whereas adoption of the sws1 cone versus rod fate during embryonic development is largely dependent on *tbx2b*; *tbx2b* is not sufficient to drive sws1 cone fate or sws1 opsin. By targeting *thrβ2* or *nrl* on our *tbx2a* and *tbx2b* single and double mutant lines, we show that both paralogues are necessary for sws1 cone specification. Secondly, we uncovered that the early differentiating photoreceptor progenitors have the capacity to adopt the lws cone, sws1 cone or rod phenotypes, whereas the potential of the late progenitor is limited to rods or sws1 cones. In juvenile and adult zebrafish, sws1 cone specification in the dorsal retina requires *tbx2b*, however both *tbx2a* and *tbx2b* are necessary for sws1 cone specification in the central and ventral retina. And following light damage in the adult, *tbx2a* haploinsufficiency markedly reduced sws1 cone regeneration. Our data reveal that in the cone-dominated zebrafish retina, *tbx2a* and *tbx2b* function to maintain the sws1 cone by repressing the rod fate in a common progenitor and provides a model for the role of transcriptional repressors in regulating the timing of photoreceptor cell fate specification from multipotent progenitors.

## Results

### *tbx2a* and *tbx2b* are necessary for sws1 cone specification from embryonic progenitors

We took advantage of the hypomorphic allele of *tbx2b*, *tbx2b^p25bbtl^ (lots-of-rods, tbx2b^lor^*) to investigate the role of *tbx2a* and putative genetic interactions of *tbx2a* and *tbx2b* across life history (Alvarez-Delfin et al. 2009). One-cell stage Tg(−5.5sws1:EGFP) and *tbx2b^lor/lor^* embryos were injected with one or two guide CRISPR/Cas9 duplexes targeting exon 1 and/or exon 2 of *tbx2a*. In the F1 larvae, we recovered 3 large deletions disrupting the DNA binding domain (*tbx2a^fl20^, tbx2a^fl24^, tbx2a^fl25^)* and two frameshift mutations both predicted to result in a premature stop codon: a 5 bp deletion upstream of the DNA binding domain (*tbx2a^fl18^*) and a 5 bp deletion within the DNA binding domain (*tbx2a^fl19^*) (Figure 1A). Heterozygous F_1_ adults were mated and in 5-day post fertilization (dpf) F_2 l_arvae the number and types of photoreceptors were quantified by immunolabelled rods (4C12) and by sws1:EGFP expression (Takechi et al. 2003). When brought to homozygosity or as compound heterozygotes with an indel over a large deletion, *tbx2a* mutant larvae displayed a small but significant reduction in sws1 cones but no discernable rod phenotype (Figure 1C). However, the numbers of rods and sws1 cones in *tbx2a^-/-^*mutant larvae were sensitive to dosage changes at the *tbx2b* locus. Larvae with a single *tbx2b^lor^* allele on a *tbx2a* mutant background displayed a significant increase in rod number and fewer sws1 cones than *tbx2a* mutants alone. As previously described, larvae homozygous for *tbx2b^lor/lor^* possessed few sws1 cones which were randomly distributed across the retina and a significant increase in rods (Figure 1D and E). *tbx2b^lor/lor^* larvae carrying a single *tbx2a* mutant allele had a further reduction in the remaining sws1 cones when compared to *tbx2b^lor/lor^* alone (Figure 1F). Lastly, all larvae homozygous for *tbx2a^-/-^; tbx2b^lor/lor^* exhibited no detectable sws1 cones (Figure 1G). All larvae showed normal optokinetic responses (OKR) for responses to rotating stripes illuminated with white light, but *tbx2a^-/-^; tbx2b^lor/lor^* failed to survive to juvenile stages, whereas all other genetic combinations were viable. Double mutants displayed several additional novel phenotypes including a single otolith, heart defects, and altered distribution of calretinin positive cells in the taste system consistent with genetic redundancy of *tbx2a* and *tbx2b* in other organ systems (Fig. S1).

**Figure 1:**
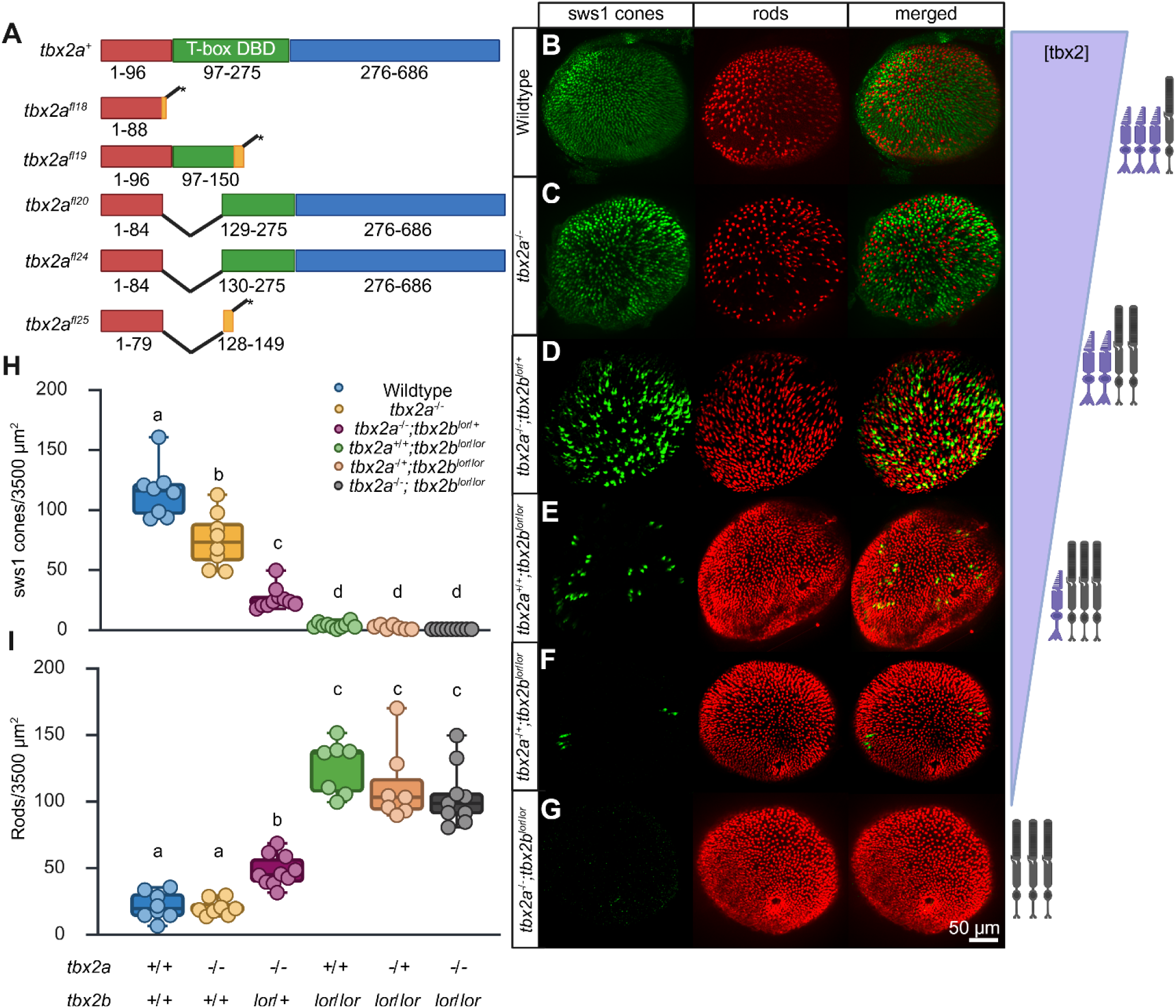
Dosage-dependent regulation of sws1 cones and rods in *tbx2a* and *tbx2b* mutant larvae. A) Diagram of wildtype Tbx2a protein illustrating the DNA binding domain (DBD) and the predicted changes in mutant protein. B-G) Wholemount immunolabeling of 5 dpf eyes for sws1EGFP (green) and rods (red). Wildtype (n=8) sws1 cone (green) and rod distribution (red) across genotypes. H) Quantification of sws1 cones in a 3500 μm^2^ area just dorsal the optic nerve: Wildtype (n=8, 114+/-22.1), *tbx2a^-/-^* (n=8, 74+/-22.7), *tbx2a^-/-^*; *tbx2b^lor/+^* (n=10, 26+/-9.3), *tbx2b^lor/lor^;tbx2a^+/+^*(n=8, 3+/-2.3), *tbx2b^lor/lor^;tbx2a^fl20/+^* (n=8, 1.6+/- 1.7), *tbx2b^lor/lor^;tbx2a^fl19/fl20^* (n=9, 0). One-way ANOVA with a Tukey’s post-hoc. p<0.0001, symbols indicate groups that are statistically different. I) Quantification of rod number in the same region; Wildtype (21+/- 10.2), *tbx2a^-/-^* (20+/-5.9), *tbx2a^-/-^*; *tbx2b^lor/+^* (47+/-11.5), *tbx2b^lor/lor^;tbx2a^+/+^* (125+/-20.0), *tbx2b^lor/lor^;tbx2a^fl20/+^* (111+/-28.6), *tbx2b^lor/lor^;tbx2a^fl19/fl20^* (104+/-22.8). One-way ANOVA with a Tukey’s post-hoc. p<0.035, symbols indicate groups that are statistically different.

To test if the *tbx2* paralogues are necessary for specification of the sws1 cone we challenged the photoreceptor progenitors by genetically manipulating *thrβ2* and *nrl* on wildtype and *tbx2* mutant backgrounds. *thrβ2* expression is necessary to specify the lws cone fate (Suzuki et al. 2013; Deveau et al. 2020; Volkov et al. 2020). Gene knockdown or genetic mutations of *thrβ2* resulted in an absence of lws cones and an increase in sws1 cones, consistent with a cell fate switch (Suzuki et al. 2013; Deveau et al. 2020; Volkov et al. 2020). If *tbx2*s are necessary to specify the sws1 cone fate, then we anticipated that no sws1 cones would be observed after targeting *thrβ2* on the *tbx2a^-/-^; tbx2b^lor/lor^* background. In preliminary studies, morphilino knockdown of *thrβ2* on the *tbx2b^lor/lor^* or *tbx2b^fby/fby^*, a putative null allele, embryos resulted in a significant reduction in arrestin3a (zpr1) immunolabelling which labels lws and rh2 cones (Renninger et al. 2011). This decrease was accompanied by a significant increase in sws1 cones, and unexpectedly, a significant increase in rod number compared to the mutant *tbx2b* phenotype alone (Fig S2). These data suggest two hypotheses; first, the increase in sws1 cones on *tbx2b* mutant background may be dependent upon expression of *tbx2a*. Secondly, the increase in rods suggests that the early photoreceptor prognitor has the potential to adopt an lws cone, sws1 cone, or rod fate. To further test progenitor fate, one cell stage embryos from mating either wildtype, *tbx2b^lor^*, or *tbx2a^fl18/fl20^*; *tbx2b^lor/+^* adults were injected with two gRNAs targeting the first exon of *thrβ2*. CRISPant larvae were immunolabelled for arrestin3a, rods and sws1:EGFP followed by PCR amplification and sequenced to detect lesions in *thrβ2*. Targeting *thrβ2* in wildtype embryos resulted in a significant decrease in arrestin3a labelling and an increase in sws1 cones but no change in rod number, as anticipated (Fig. S2, Figure 2B-B’) (Suzuki et al. 2013; Deveau et al. 2020; Volkov et al. 2020). *tbx2b^lor/lor^; CRISPant thrβ2* animals displayed a significant increase in both sws1 cones and rods compared to *tbx2b^lor/lor^* alone much like the *thrβ2* morpholino injected embryos (Figure 2D-D’). However, when targeting *thrβ2* on *tbx2a^fl19/fl20^; tbx2b^lor/lor^*embryos, no sws1 cones were observed, and the number of rods was significantly increased over the *tbx2a^-/-^; tbx2b^lor/lor^*mutant alone (Figure 2F-F’). These data are consistent with the hypotheses that the *tbx2*s are necessary for sws1 cone specification and that the increase in the number of sws1 cones following knockdown of *thrβ2* on *tbx2b^lor^* and *tbx2b^fby^* is dependent upon *tbx2a* expression. Additionally, the significant increase in rods when *thrβ2* and *tbx2*s are manipulated is evidence that the early photoreceptor progenitor is multipotent with the potential to adopt the lws cone, sws1 cone or rod fate.

**Figure 2:**
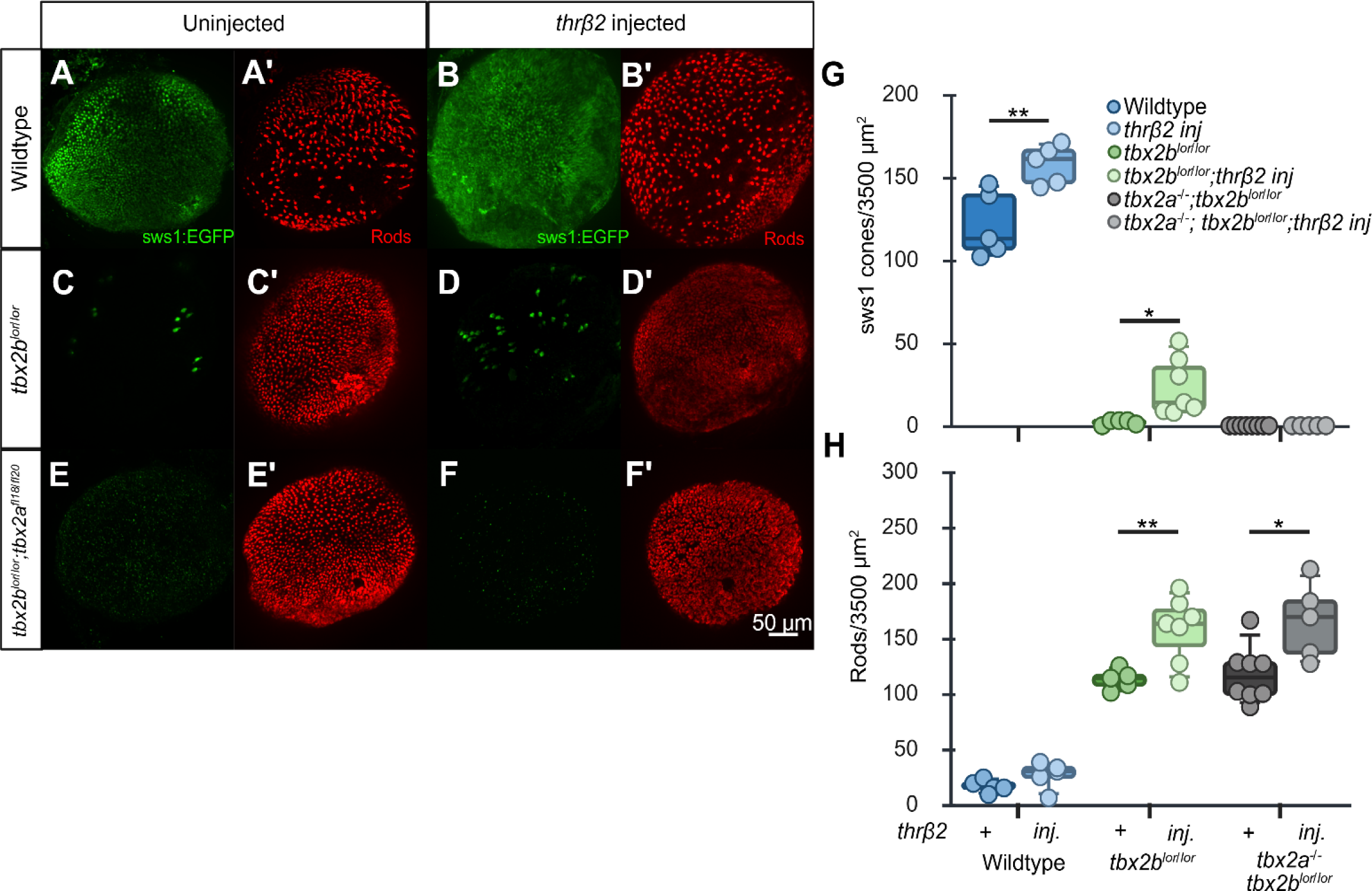
*tbx2*s are necessary for sws1 cone fate specification. A-F) Wholemount immunolabeling of 5 dpf eyes for sws1EGFP (green A-F) and rods (A’-F’) across genotypes. G) Quantification of sws1 cones in a 3500μm^2^ square dorsal the optic nerve opposite the ventral patch phenotypes: Wildtype (n=5, 121+/-19.8), *thrβ2* inj. (n=5, 158+/-11.8), *tbx2b^lor/lor^* (n=5, 2+/-1.4), *tbx2b^lor/lor^ thrβ2* inj. (n=7, 23+/-17*), tbx2a^-/-^; tbx2b^lor/lor^* (n=8, 0), *tbx2a^-/-^; tbx2b^lor/lor^ thrβ2* inj (n=5, 0). H) Quantification of rod number in the same region: Wildtype (16+/-5.5), *thrβ2* inj. (26+/-12.3), *tbx2b^lor/lor^* (113+/-9.0), *tbx2b^lor/lor^ thrβ2* inj. (158+/-29.7), *tbx2a^-/-^; tbx2b^lor/lor^* (117+/-25), *tbx2a^-/-^; tbx2b^lor/lor^ thrβ2* inj. (162+/-32.2). Student’s unpaired t-test, * p<0.05, **p<0.01.

Photoreceptor specification is temporally regulated; the majority of cones are derived from progenitors that exit the cell cycle early in embryonic neurogenesis, whereas rods are derived from progenitors exiting the cell cycle later in neurogenesis (Schmitt and Dowling, 1999; Cepko et al. 1996). We tested if *tbx2a* and *tbx2b* are necessary for respecification of rod photoreceptor progenitors into sws1 cones. Across species, mutations of *nrl* result in a loss of rods and an increase in sws1 cones; And *nrl^-/-^; tbx2b^lor/lor^* also displayed an increase in sws1 cones (Mears et al. 2001; Oh et al. 2007; Oh et al. 2008; Oel et al. 2020; Liu et al. 2022; Neil et al. 2024). If *tbx2a* and *tbx2b* are necessary for sws1 cone specification then we anticipate that targeting *nrl* on the *tbx2a^-/-^; tbx2b^lor/lor^* background would result in a decrease in rods but no increase in sws1 cones. *tbx2a^fl18/fl20^;tbx2b^lor/+^* animals were inbred, and progeny injected with 3 gRNAs targeting *nrl*. *nrl* CRISPant wildtype larvae displayed a significant reduction in rods, as expected (Figure 3B-B’). Targeting *nrl* on *tbx2b^lor^*had a significant reduction in rods accompanied with a slight, albeit significant, increase in sws1 cones (Figure 3D-D’). Surprisingly, the distribution of sws1 cones within the retina was limited to the equitorial zone, which corresponds to the visual streak in larval zebrafish (Figure 3D) (Yoshimatsu et al. 2020). In contrast, no sws1 cones were observed following targeting of *nrl* on the *tbx2a^fl18/fl20^*;*tbx2b^lor/lor^*, confirming that *tbx2a* and *tbx2b* are necessary for sws1 cone specfication (Figure 3F-F’). However, we did not observe an increase in arrestin3a or sws2 cone labeling suggesting that late stage progenitors were fate restricted.

**Figure 3:**
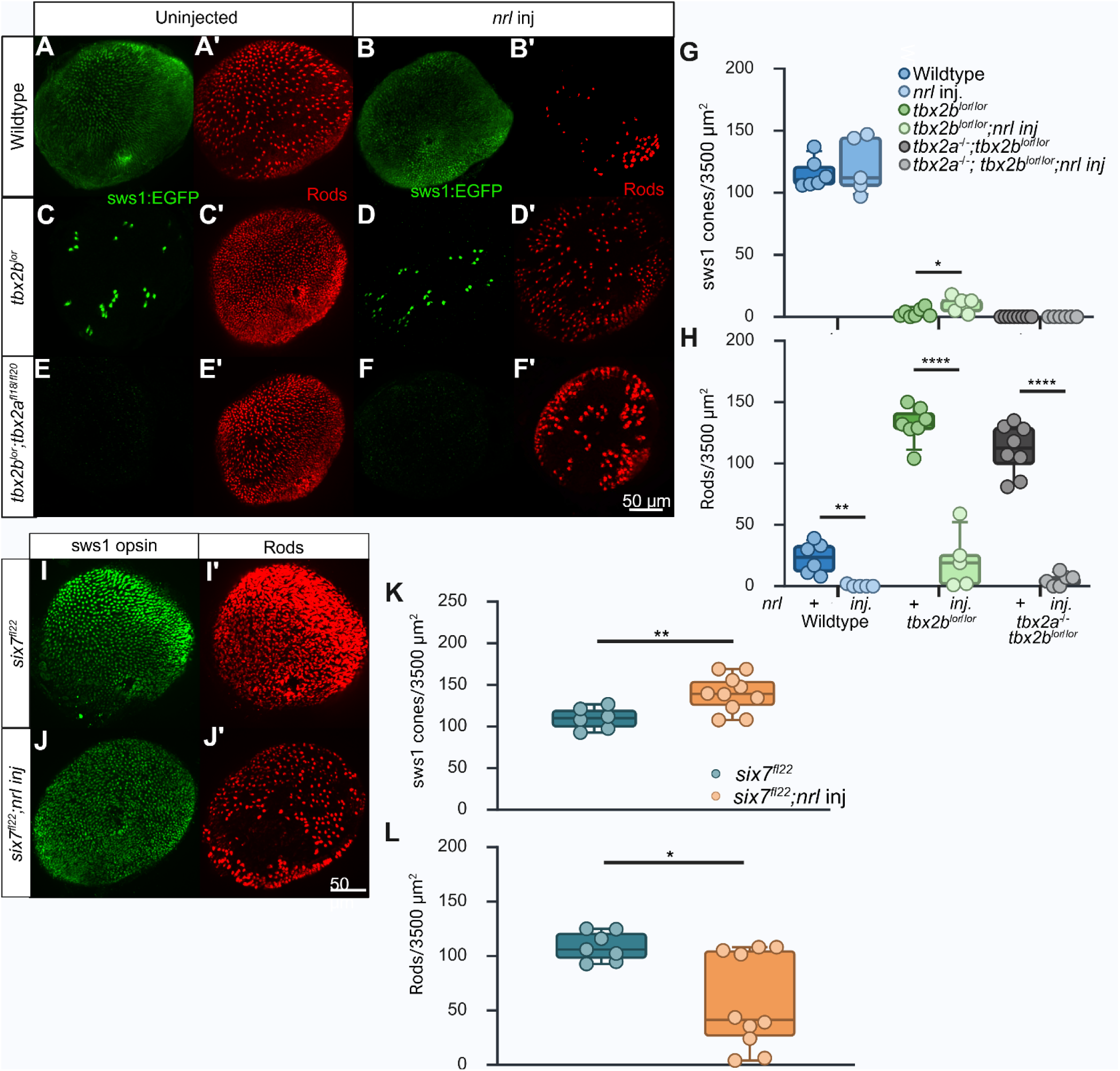
The late photoreceptor progenitor is fate restricted. . A-F) Wholemount immunolabeling of 5 dpf eyes for sws1EGFP (green A-F) and rods (A’-F’) across genotypes. G) Quantification of sws1 cones in a 3500μm^2^ square dorsal the optic nerve opposite the ventral patch phenotypes: Wildtype (n=6, 121+/- 24.1) *nrl* inj. (n=5, 121+/-22.8), *tbx2b^lor/lor^*(n=7, 3+/-2.3), *tbx2b^lor/lor^* nrl inj. (n=5, 10+/-6.5), *tbx2a^-/-^; tbx2b^lor/lor^* (n=8, 0) *tbx2a^-/-^; tbx2b^lor/lor^ nrl* inj. (n=7, 0). H) Quantification of rods in the same region: Wildtype (23+/- 12.7) *nrl* inj. (1+/-0.89), *tbx2b^lor/lor^* (132+/-14.8), *tbx2b^lor/lor^* nrl inj. (21+/-23.6), *tbx2a^-/-^; tbx2b^lor/lor^* (111+/-20.4) *tbx2a^-/-^; tbx2b^lor/lor^ nrl* inj. (5+/-5.0). Student’s unpaired t-test, * p<0.05, **p<0.01, **** p<0.0001. I-J) Wholemount immunolabeling of 5 dpf eyes for sws1EGFP (green I,J) and rods (I’,J’) across mutants. K) Quantification of sws1 cones in a 3500μm^2^ square dorsal the optic nerve opposite the ventral patch phenotypes: *six7^fl22/fl22^*(n=7, 110+/-13.1), *six7^fl22/fl22^; nrl* inj. (n=10, 139+/-21.9). Student’s unpaired t-test, ** p<0.01. L) Quantification of rod number in the same region: *six7^fl22/fl22^* (109+/-13.4)*, six7^fl22/fl22^*; *nrl* inj. (58+/-43.4). Mann-Whitney test, *p<0.05.

To expand these findings that late progenitors can be respecified as sws1 cones, we tested the effect of targeting *nrl* on the *six7* mutant embryos. *six7* regulates the number of rods in larval zebrafish retinas by repressing the mitosis of the late stage progenitors (Sotolongo-Lopez et al. 2016; Ogawa et al. 2015; Ogawa et al. 2019). Knockdown or genetic mutations of *six7* result in a significant increase in rods associated with extended mitosis of retinal progenitors with the majority of these differentiating as rods (Sotolongo-Lopez et al. 2016; Ogawa et al. 2015; Ogawa et al. 2019; Saade et al. 2013). A novel allele of *six7,* was generated using CRISPR/Cas9 duplex targeting of exon 1 of *six7*. An allele bearing a 1 bp deletion (*six7^fl22^*) resulting in a predicted premature stop codon within the Six7 homeodomain was recovered in the F1 generation.

Homozygous *six7^fl22^* larvae demonstrate a significant increase in rods and lack of green photoreceptors as previously described (Sotolongo-Lopez et al. 2016; Ogawa et al. 2015; Ogawa et al. 2019). Targeting of *nrl* on *six7^fl22/fl22^* resulted in a significant reduction in rods accompanied by a significant increase in sws1 cones (Figure 3L). Together, these data suggest that late stage progenitors retain potential to differentiate into rods as well as sws1 cones.

### *tbx2a* and *tbx2b* are necessary for sws1 cone differentiation from the ciliary marginal zone

Having established that *tbx2a* and *tbx2b* are necessary for specification of sws1 cone versus rod fate in embrynoic neurogenesis, we next sought to test their roles in juvenile and adult neurogensis. Throughout life, teleosts, including zebrafish, show continuous neurogenesis at the retinal margin from populations of mitotic stem cells located at the CMZ (Raymond et al. 2006). Analagous to rings on a tree, older tissue is nearer the central retina and younger tissue nearer the margins. To demarkate retinal outgrowth from the CMZ, mitotic cells and their descendants were labeled with EdU at 30 dpf or 30dpf and again at 60 dpf (Figure 4A-D). Histological sections were taken along the dorsal-ventral axis through the optic nerve.

**Figure 4:**
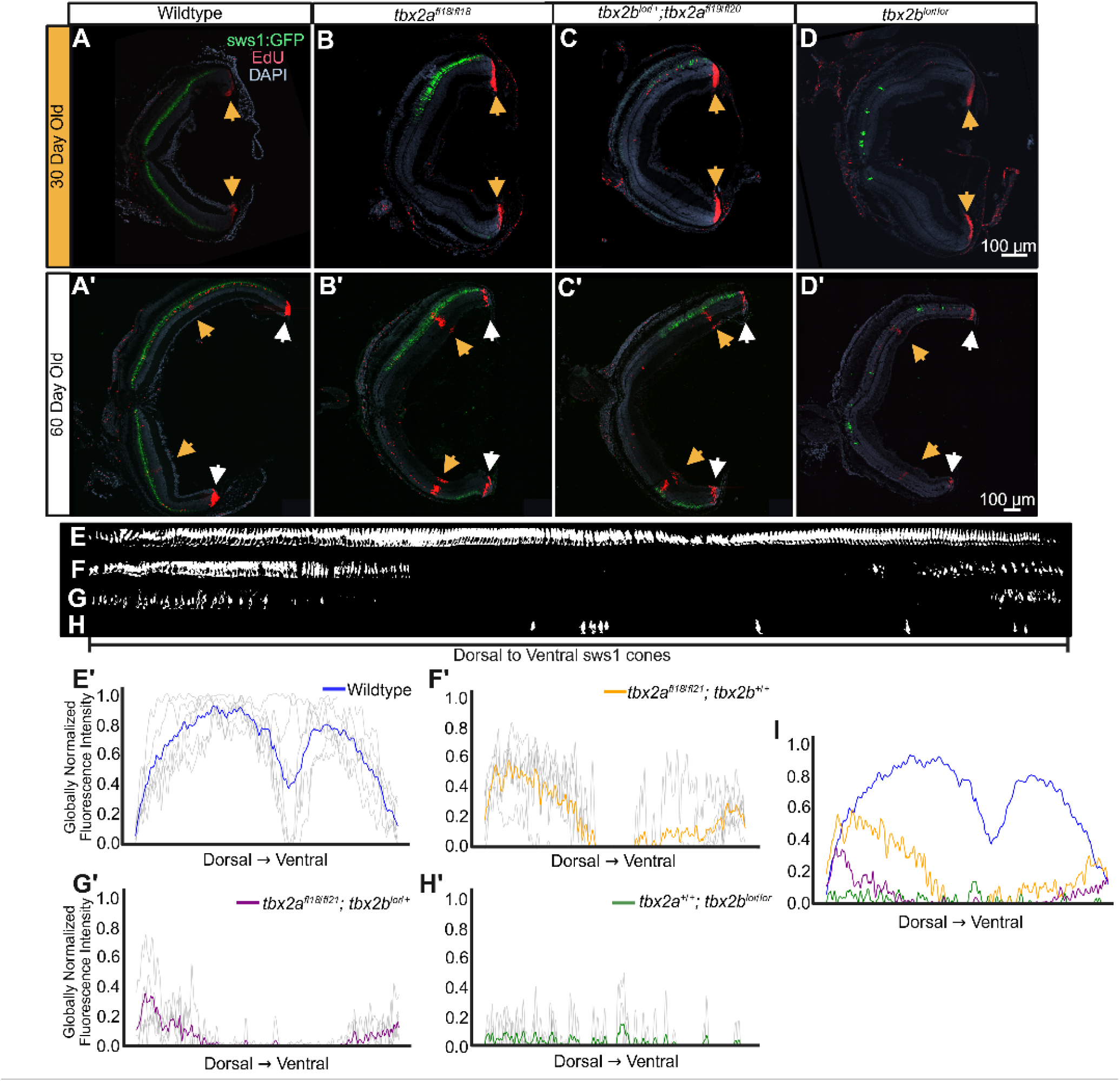
*tbx2a* and *tbx2b* demonstrate epistasis in sws1 cone differentiation from CMZ derived progenitors. A-D) 30 dpf retinal cross section immunolabeling for sws1 cone (green) and EdU (red) distribution across genotypes. Yellow arrowhead represents the 30 day EdU treatment. (A’-D’) 60 dpf retinal cross section immunolabeling for sws1 cones (green) and EdU (red) across genotypes. Yellow arrowhead demarcate the 30 day old EdU treatment and the white arrowhead marks the 60 dpf EdU injection. E-H) 60 dpf sws1 cones linearized from dorsal to ventral retina and thresholded across genotypes. E’-H’) Globally normalized sws1 cone distributions from dorsal to ventral retina across genotypes. I) Summary plot of sws1 distribution for all genotypes at 60 dpf: Wildtype (n= 6), tbx2a^-/-^ (n=5) tbx2a^-/-^;tbx2b^lor/+^ (n=5), tbx2b^lor/lor^ (n=5).

At 30 dpf, the majority of the EdU labelling was localized to the mitotic cells of the CMZ in the dorsal and ventral retina, and rod progenitors scattered along the ONL. At 60 dpf, two bands of EdU labeling were present: the CMZ in the dorsal and ventral retina as well as bands of neurons closer to the central retina, which represent the cells that exited the cell cycle at 30 dpf post EdU uptake (Fig 4 A’-D’). In 60 dpf fish, the relative expression of EGFP in the Tg(−5.5sws1:EGFP) was quantified as a surrogate for the density of sws1 cones (Fig. 4E-H). In wildtype animals at 30 and 60 dpf, sws1 cones formed a nearly uniform row in the outer nuclear layer (ONL) running from the dorsal to ventral retina. Flourescent intensity tapered near the retinal margins, and was absent at the optic nerve head where no photoreceptors are present (Figure 4E-E’). *tbx2a^fl19/fl20^*animals displayed a distinct dorsal-ventral asymmetry in the distribution of sws1 cones. The flourescent intensity was highest in the dorsal retina near the CMZ, with a clear absence of sws1 cones in a zone stretching from dorsal to the optic nerve to the ventral margin. sws1 cones were observed in the retinal outgrowth from 30 to 60 dpf with fewer and sparse labeling at the ventral margin (Figure 4F-F’). A single *tbx2b^lor^*allele on the *tbx2a^fl19/fl20^* resulted in an expanded region with no EGFP expression and sparse labeling in both the dorsal and ventral margins at 60 dpf (Figure 4G-G’). Simliar to our previous reports, sws1 cones were virtually absent in *tbx2b^lor^*animals, but when present, were sporadically distributed across the retina including the central retina (Figure 4H-H’).

*tbx2a* and *tbx2b* are expressed by mature sws1 cones (Ogawa et al. 2021; Ogawa and Corbo, 2021; Angueyra et al. 2023; Farrell et al. 2018; Sur et al. 2023). TUNEL was used to test if *tbx2a* is necessary for survival of sws1 cones. If so, we anticipated observing increased cell death near the margins. However, no difference in the number of TUNEL positive cells was observed across genotypes (Fig S3). Taken together, these data suggest that *tbx2a* and *tbx2b* have overlapping yet distinct spatial and temporal requirements during outgrowth from the CMZ, and as in larvae, *tbx2a^-/-^*fish were sensitive to gene dosage at the *tbx2b* allele.

### *tbx2b* is not sufficient for sws1 cone specification

To determine if *tbx2b* is sufficient to drive sws1 opsin expression or sws1 cone fate, an EGFP:tbx2b fusion protein was overexpressed in post-mitotic cones in transgenic larvae downstream of the *gnat2* promoter (Kennedy et al. 2007). Overexpression of *thrβ2* using the *gnat2* led to an increased number of lws cones and lws opsin expression in other cone subtypes (Suzuki et al. 2013). A transgenic line was established by injection of single-cell stage wildtype embryos with transposase mRNA and a plasmid containing a *(Tol2-gnat2:EGFP-tbx2b-polyA)* cassette and a *cmlc2:EGFP* cassette that drives heart specific expression of EGFP (Kwan et al. 2007). Embryos were grown to adults and mated. F_1_ embryos were screened for cardiac expression of EGFP. Immunohistochemistry was performed at 6-dpf to determine the fate of EGFP expressing cone cells.

In transgenic larvae, *EGFP-tbx2b* expression was localized to the outer nuclear layer of the retina. In transgenic animals with lower levels of *EGFPtbx2b* expression, the EGFP was restricted to the majority of cone nuclei. High expression of *EGFPtbx2b* resulted in EGFP fluorescence throughout the cone cell body and terminal, yet in no case was the number of cells immunolabelled for sws1 opsin changed from wildtype, nor was there an increase in sws1 opsin expression by qPCR (Figure 5E). These data suggest that *tbx2b* alone is not sufficient to drive sws1 cone fate or sws1 opsin in larval zebrafish photoreceptors. Transgenic larvae were co-immunolabeled with a monoclonal antibody, zpr3, which recognizes rhodopsin (rho) and rh2, or arrestin3a and a polyclonal antibody to sws2 opsin (Hu et al. 2024). Overexpression of *tbx2b* resulted in reduced immunolabelling for arrestin3a, rh2, and sws2 opsin and the significant reduction in sws2, rh-2, and lws opsin expression by qPCR, but no discernable change in cell number (Figure 5A-E). These data support the hypothesis that *tbx2b* acts as a repressor (Alvarez-Delfin et al. 2009; Zhu et al. 2014; Cho et al. 2017; Teegala et al. 2018).

**Figure 5:**
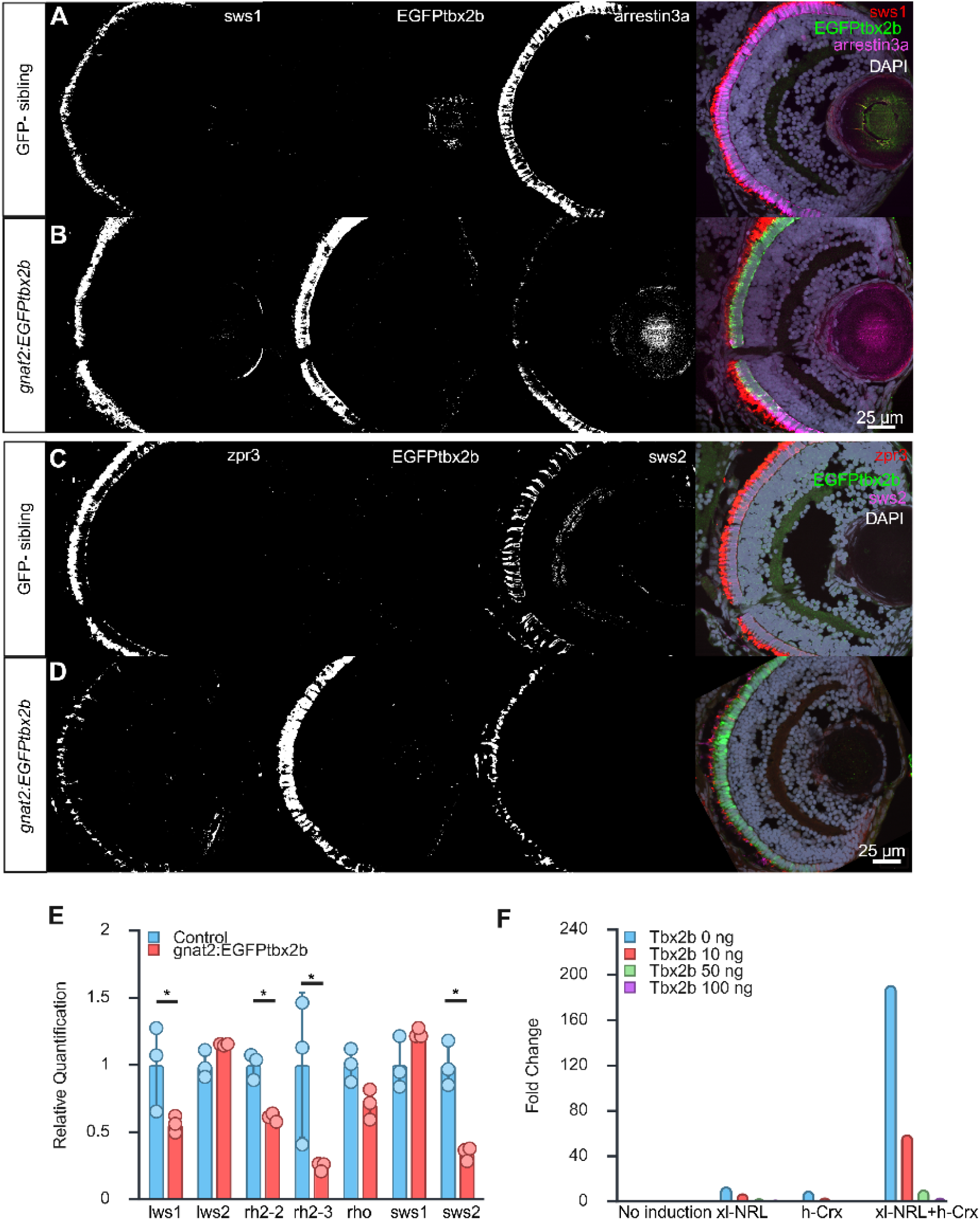
Overexpression of *tbx2b* was not sufficient to drive sws1 opsin expression. A-B) 6-dpf retinal cross sections immunolabelled for sws1 opsin (red), EGFPtbx2b (green), and arrestin3a (magenta) in wildtype and transgenic larvae. C-D) 6 dpf retinal cross sections immunolabelled for zpr3 opsin (red), EGFPtbx2b (green), and sws2 (magenta) in wildtype and transgenic larvae. E) qPCR for cone opsin subtype and rhodopsin. Mann-Whitney one-way t-test p=0.05. (n=3) F) Fold change of rho:luciferase demonstrates dosage-dependent repression at the rho promoter (n=3).

The mechanism leading to a cell fate switch of sws1 cone precursors into rods in *tbx2a* and *tbx2b* mutants remained as an outstanding question. To address this, a promoter reporter assay using human-CRX, *Xenopus*-Nrl, zebrafish-Tbx2b, and *Xenopus-(−*5.5Rho:luciferase) was performed in a heterologous expression system. HEK-293 cells were co-transfected with constructs containing human-CRX or *Xenopus*-Nrl and *Xenopus*-(−5.5Rho:Luciferase). h-CRX or X-Nrl alone activated transcription at the rhodopsin promoter approximately 10-and 13-fold respectively. However, co-transfection of h-CRX and X-Nrl synergistically drove a 200-fold increase in luciferase expression from the 5.5 kb rhodopsin promoter, as expected (Chen et al. 1997; Whitaker and Knox, 2004) (Figure 5F). The addition of a construct containing z-Tbx2b significantly inhibited the synergistic activity of h-CRX and X-Nrl. Additionally, increasing concentrations of z-Tbx2b resulted in a dosage-dependent decline in luciferase activity to near uninduced levels at 100 ng z-Tbx2b vector (Figure 5F).

### *tbx2a* is necessary for sws1 cone regeneration following light damage

Zebrafish possess the capacity for regeneration of neurons including photoreceptors from Muller glia derived progenitors (Lenkowski and Raymond, 2014; Cameron and Easter, 1995; Vhitelic and Hyde, 2000; McGinn et al. 2018). To test the roles of *tbx2a* and *tbx2b* during regeneration, >6 month old, age and size matched *Tg(−5.5sws1:EGFP);* wildtype, heterozygous and homozygous *tbx2b^lor^*and *tbx2a^fl18^* adult zebrafish underwent high intensity light damage (>120,000 lux) followed by 96 hours of continuous light (20,000-60,000 lux) and 10 days recovery, as previously described (Figure 6A) (Thomas and Thummel, 2013).

**Figure 6:**
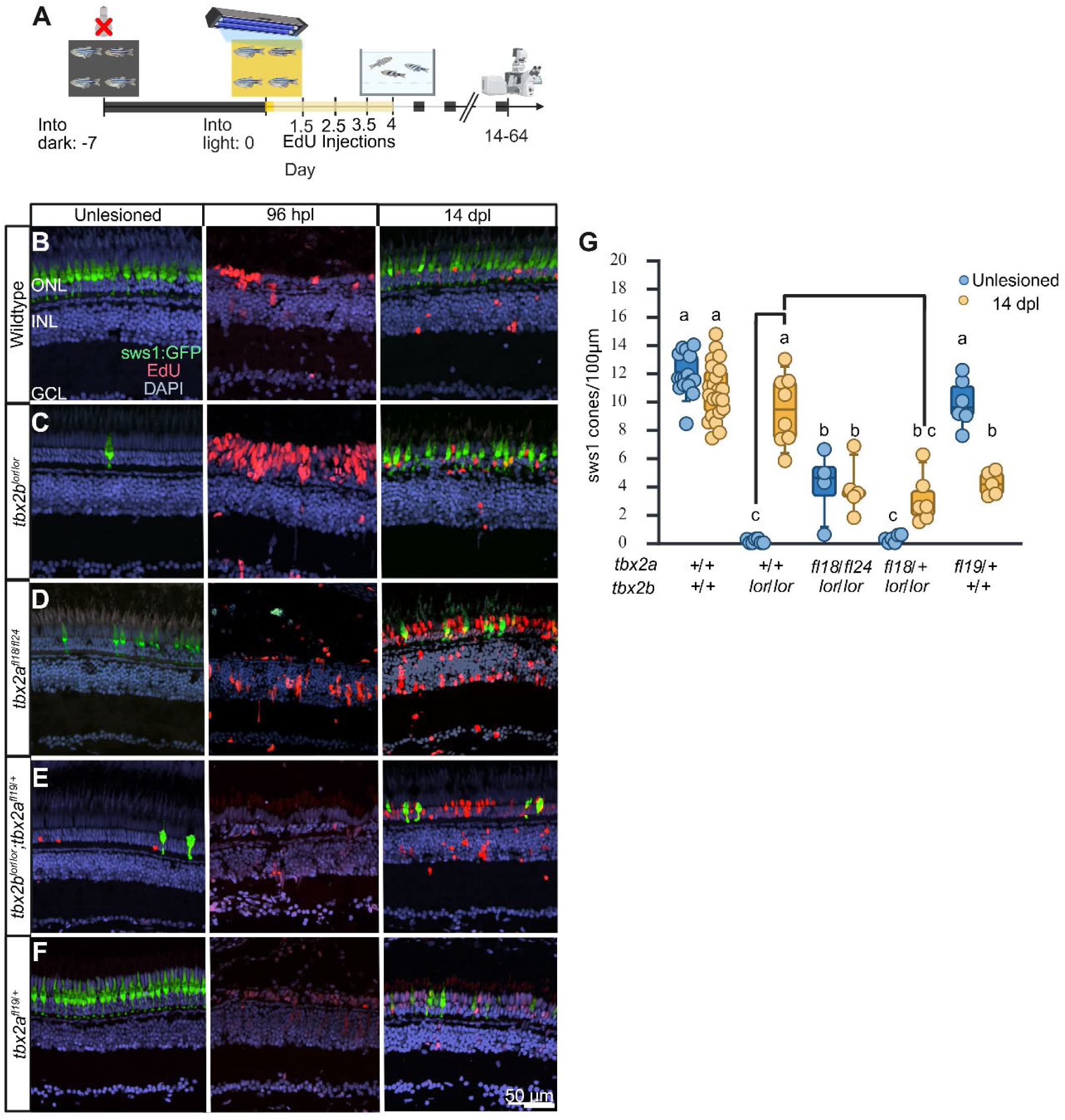
*tbx2a* is required for sws1 cone differentiation from Muller glia derived progenitors. A) Schematic of light damage protocol timeline with day 0 being the onset of light damage. B-F) Uninjured adult retinal cross sections immunolabeled for sws1 cones (green) and EdU (red) just dorsal the optic nerve across genotypes (Left column). 96 hours post light damage (hpl) severely disrupted the outer nuclear layer (ONL) and increased labelling for EdU (Center). 14 days post light damage (dpl) immunolabeling for sws1 cones (green) and EdU (red) across genotypes (right column). number of sws1 cones. G) Quantification of sws1 cones per 100 μm in damaged (blue) and 14 dpl (yellow) across genotypes: Wildtype unlesioned (n=16, 12+/-1.4), wildtype 14 dpl (n=18, 11+/-2.0), *tbx2b^lor/lor^*unlesioned (n=7, 0.1+/-0.1), *tbx2b^lor/lor^* 14dpl (n=8, 9+/-2.5), *tbx2a^fl18/fl24^*unlesioned (n=4, 4+/-2.5), *tbx2a^fl18/fl24^* 14 dpl (n=5, 4+/-1.9), *tbx2b^lor/lor^;tbx2a^fl18/+^* unlesioned (n=6, 0.3+/-1.8), *tbx2b^lor/lor^;tbx2a^fl18/+^* 14dpl (n=6, 3+/-1.8), *tbx2a^fl19/+^* unlesioned (n=6, 10+/-1.7), *tbx2a^fl19/+^* 14dpl (n=5, 4+/- 0.8). Two-way ANOVA with Bonferroni post-hoc, symbols indicate significantly different groups, p< 0.0001.

In the central retina of undamaged wildtype fish, the pattern of GFP expression was consistent with 60 dpf zebrafish. The sws1 cone cell bodies and outer segments formed a uniform row in the middle of the outer nuclear layer, with rod nuclei more vitreal and long cone nuclei more distal. Few, if any sws1, cones were observed in undamaged *tbx2b^lor/lor^* and *tbx2b^lor/lor^; tbx2a^fl20/+^*animals (Figure 6B, C, E), whereas *tbx2a^-/-^*animals possessed an intermediate number of sws1 cones (Figure 6D). At 96 hours post light onset, all treatment groups showed thinning of the outer nuclear layer in the central retina and no GFP positive cells remained (Figure 6B-E). EdU positive cells were observed in the inner and outer nuclear layers consistent with proliferating Muller glia derived progenitors (Campbell et al. 2021; Campbell et al. 2023). 14 days post lesion (dpl), EdU positive cells were found within the outer nuclear layer consistent with photoreceptor regeneration.

The number of sws1 cones in regenerated retinas varied across genotypes. The number of sws1 cones was counted in the central retina, dorsal to the optic nerve. Following recovery in light-damaged wildtype zebrafish, the number of regenerated sws1 cones did not significantly differ from undamaged siblings (Figure 6B). Unexpectedly, following recovery, *tbx2b^lor/lor^* retinas possessed a far greater number of sws1 cones than undamaged siblings. In fact, the number of regenerated sws1 cones was no different than that observed in wildtype animals (Figure 6C). These data suggest that *tbx2b* is not necessary for regeneration of sws1 cones. Conversely, following light-damage of *tbx2a^fl18/fl24^* adults the number of sws1 cones was equal to that of the undamaged siblings but that number was significantly fewer than was observed in wildtype or *tbx2b^lor/lor^*animals (Figure 6C). To test if *tbx2a* is responsible for regeneration of sws1 cones in *tbx2b^lor/lor^*, a single *tbx2a^fl18^* allele was placed on the *tbx2b^lor/lor^* background. Following light damage and recovery, the number of sws1 cones in *tbx2b^lor/lor^; tbx2a^fl18/+^* animals was greatly reduced compared to *tbx2b^lor/lor^* alone (Figure 6E). The reduction in sws1 cone regeneration in *tbx2b^lor/lor^; tbx2a^fl18/+^* opened the possibility that sws1 cone regeneration requires two functional alleles of *tbx2a*. To test this, we light damaged heterozygous *tbx2a^fl18/+^* retinas. The number of sws1 cones in undamaged *tbx2a^fl18/+^* was no different than undamaged wildtype controls (Figure 6G). However, following light damage and recovery, *tbx2a^fl18/+^* adults showed significantly reduced sws1 cone regeneration similar to those of *tbx2b^lor/lor^; tbx2a^fl18/+^* and *tbx2a^fl18/fl24^* animals (Figure 6F). These data are consistent with the hypothesis that *tbx2a* is the primary driver of sws1 cone regeneration and *tbx2a* is haploinsufficient.

## Discussion

In this work, we provide several important mechanisms regulating photoreceptor specification in zebrafish. Our data show that the roles of *tbx2a* and *tbx2b* shift in photoreceptor development across ontogeny. During embryonic neurogenesis, *tbx2b* expression primarily facilitates specification of the sws1 cone versus rod fate, whereas in juvenile fish, *tbx2a* and *tbx2b* show regional differences during retinal outgrowth from the CMZ. Yet, following light damage, sws1 cone regeneration is primarily dependent on expression of *tbx2a*. Additional genetic manipulations directly support the hypothesis that early photoreceptor progenitors have the potential to adopt multiple late photoreceptor fates, but late-stage progenitors are restricted in their developmental potential. We propose that in zebrafish, the fate of the multipotent progenitors is restricted by the expression of transcriptional repressors which initially inhibit differentiation then narrow the competency to one of two alternative fates.

### *tbx2a* and *tbx2b* are necessary for sws1 cone specification

We uncovered that *tbx2* paralogues are necessary for sws1 cone specification and likely function in the same pathway to regulate sws1 cone versus rod specification. As previously reported and shown here, mutations of *tbx2b* resulted in few sws1 cones and a significant increase in rods in larvae, evidence that *tbx2b* is the primary determinant of sws1 cone versus rod fate (Alvarez-Delfin et al. 2009). Mutations of *tbx2a* resulted in only a modest reduction in sws1 cones in larvae and no discernable change in rod number. However, larvae homozygous for mutant alleles of *tbx2a* or *tbx2b* were sensitive to changes in gene dosage of the paralog such that a single *tbx2b^lor^* allele on *tbx2a* homozygous larvae resulted in a dramatic reduction in the number of sws1 cones and a significant increase in rods. Lastly, *tbx2b^lor/lor^; tbx2a^-/-^*larvae lacked sws1 cones even when challenged with additional genetic manipulations.

Targeting single genes or identification of the causative variants underlying human disease have provided a model whereby transcription factors direct the photoreceptor progenitor to adopt one of two fates (Aramaki et al. 2022). Targeting *thrβ2* or *nrl* on the *tbx2a^-/-^; tbx2b^lor/lor^* double mutants revealed genetic interactions in determining photoreceptor identity. The current model of retinal neurogenesis suggests that mitotic progenitors pass through a series of competency states (Cepko, 1996). Our data support a model in zebrafish where a photoreceptor progenitor passes through a hierarchy in fate specification. In targeting *thrβ2* on the *tbx2b^lor/lor^*, not only did sws1 cone number increase, but rod number also significantly increased. Upon targeting *thrβ2* on the *tbx2a^-/-^;tbx2b^lor/lor^* background, a significant increase in rod number was observed however no sws1 cones were detected. These data demonstrate that a common, early, *thrβ2+* progenitor has the potential to adopt a rod fate, in addition to an lws or sws1 cone fate, and suggest interplay between the expression of *tbx2* paralogues and *nrl* in regulating the ratio of sws1 cones to rods. Whether this is a direct change from an lws cone to a rod or if the cells pass through a quasi sws1 cone fate before adopting the rod fate is unknown from the data. In mouse, *nrl* and *nuclear receptor subfamily 2 group e member 3* (*nr2e3)* repress the sws1 cone fate, and mutations of *nr2e3* result in cells adopting mixed rod/cone phenotype (Cheng et al. 2006; Cheng et al. 2006). Subsequent data revealed that the early photoreceptor progenitors that generate rods in mice temporarily activate the sws1 promoter, although this was not detected in wildtype larval zebrafish (Cheng et al. 2006; Qin et al. 2009). These data open the possibility that the function of *tbx2* paralogues in cone-dominated retinas is analogous yet opposite to the role of *nr2e3* in rod-dominated retinas.

Targeting *nrl* on *six7^fl22/fl22^* demonstrated that late-stage proliferating rod progenitors have the potential to adopt an sws1 cone fate. But no sws1 cones were observed upon targeting *nrl* on the *tbx2a^-/-^;tbx2b^lor/lor^*background, confirming the necessity of *tbx2a* and *tbx2b* for sws1 cone specification. At this time, the fate of these photoreceptor progenitors remains unknown. We observed no increase in markers of lws/rh2 cones or sws2 opsin. Together, these data are consistent with the potential of late-stage photoreceptor progenitors being restricted. Recent data in *Drosophila* have identified key regulators for the transition from early to late neural progenitors (Morales et al. 2026; Shen et al. 2026). These changes in progenitor competency from early to late cell fates could be driven by chromatin remodeling (Aramaki et al. 2022; Lyu et al. 2021).

Based upon these data, we propose that *tbx2b* acts as a repressor during larval photoreceptor cell development to antagonize cells adopting a rod fate. In zebrafish, mitotic progenitors express *crx*, *nrl*, *nr2e3*, *thrβ2*, and *tbx2a* and *tbx2b* (Ogawa and Corbo, 2021; Angeuyra et al. 2023; Farrell et al. 2018, Ogawa et al. 2021; Sur et al. 2023; Chen et al. 2005; Morris et al. 2008; Nelson et al. 2008). Yet a cell adopts a single, clearly defined cell fate. The absence of *tbx2* expression derepresses *crx* and *nrl*, driving the progenitor to a rod fate. In vitro promoter-reporter assays demonstrate Tbx2b represses the synergistic activity of Crx and Nrl at the *rhodopsin* promoter in a dosage-dependent manner. In line with these data, overexpression of *Tbx3* in mice led to a reduction in rods (Lyu et al. 2021). We propose *tbx2*s repress the rod fate, and a yet unidentified factor is necessary to drive the sws1 cone fate. Furthermore, overexpression of Tbx2b in cone precursors was not sufficient to drive the sws1 cone fate or sws1 opsin expression in other cone subtypes. In fact, overexpression of Tbx2b resulted in repression of the other cone opsins, though cell number remained unchanged. For comparison, overexpression of *thrβ2* using same promoter was sufficient for increased lws opsin expression and mixed cone fates (Suzuki et al. 2013). Correspondingly, knockdown of *tbx2a* or *tbx2b* resulted in misexpression of rh2 opsin in lws and sws2 cones respectively, supporting a role for *tbx2*s to repress misexpression of other cone opsins (Angueyra et al. 2023). Similar roles on opsin expression were observed in other fishes. *tbx2a* functionally diverged in different species of African cichlids (Sandkam et al. 2020). A shift in spectral sensitivity in some species of cichlids from rh2 to lws opsin was correlated with *tbx2a* expression differences associated with changes in cis-regulatory activity (Sandkam et al. 2020). Overall, the genetic data and the expression of *tbx2* orthologues in multiple cell types support their function as repressors.

### Roles *of tbx2a* and *tbx2b* shift throughout ontogeny

The continued neurogenesis at the retinal margin in aquatic vertebrates provides an opportunity to test the role of transcription factors in post-embryonic development. Progenitors at the CMZ recapitulate expression of many developmentally regulated transcription factors (Stenkamp et al. 1997; Perron et al. 1998). The sporadic distribution sws1 cones in *tbx2b^lor/lor^* juvenile and adult fish demonstrates that *tbx2b* remains the primary regulator for sws1 cone fate. The lack of sws1 cones directly dorsal to the optic nerve and in the ventral retina at 30 dpf in *tbx2a* mutants establishes that the initial outgrowth from the dorsal and ventral CMZ requires *tbx2a,* and may predict a potential role in dorsal-ventral patterning of the retina (Yoshimatsu et al. 2020; Ayten et al. 2025). The necessity of a functional allele at each locus for sws1 cone specfification in the dorsal and ventral retina in juvenile zebrafish demonstrates a classic example of reciprocal recessive epistasis. The further reduction in sws1 cones in the dorsal retina and ventral margin of *tbx2a^-/-^; tbx2b^lor/+^*animals demonstrates that *tbx2a* masks a dosage-dependent requirement for *tbx2b* as retinal outgrowth continues. In humans, heterozygous frameshift mutations of *TBX2* have been associated with hearing loss and incompletely penetrant nystagmous (Hua et al. 2025). Furthermore, mutations of other t-box family transcription factors present more moderate disease phenotypes in individuals who are heterozygous rather than homozygous, many of which are lethal (Papaioannou, 2014). Together, these findings suggest some level of selective pressure in zebrafish to maintain both loci.

Previous studies have shown that following acute or chronic damage, genes associated with retinal development were upregulated (Thummel et al. 2008; Morris et al. 2011; Saade et al. 2013), yet few studies have used genetic mutants to test the role of developmentally regulated genes in photoreceptor regeneration (Qin et al. 2009). Surprisingly, following light damage, *tbx2b^lor/lor^* animals regenerated sws1 cones to a wildtype level. *tbx2b^lor^* is a hypomorphic allele but the molecular lesion has yet to be identified. Our original hypothesis was that the unidentified lesion could affect a cis-regulatory element governing *tbx2b* expression during embryonic development but is not needed during regeneration. However, animals harboring a single *tbx2a* mutant allele on the *tbx2b^lor/lor^* background regenerated far fewer sws1 cones. In fact, retinas of animals heterozygous for a single *tbx2a* mutant allele which initially display a wildtype number of cones were haploinsufficient for sws1 cone following light damage. These data are consistent with subfunctionalization of *tbx2* paralogues. *tbx2b* primarily functions in scheduled sws1 cone specification during embryonic development and outgrowth at the retinal margin, whereas *tbx2a* regulates unscheduled sws1 cone specification during regeneration.

Subfunctionalization of *tbx2* paralogues is similar to observations for *nrl* and a second MAF transcription factor in zebrafish, *mafba*. Larval rod genesis is *nrl* dependent, however in juvenile and adult stages *mafba* has been shown to drive the rod fate in the absence of *nrl*, though at a reduced level (Oel et al. 2020; Liu et al. 2022). RNA expression analysis suggests that *mafba* is expressed in progenitors that are direct descendants of Muller glia (Liu et al. 2022). These data open the possibility of coevolution of *tbx2b* and *tbx2a* with *nrl* and *mafba,* respectively. Perhaps *tbx2* expression has diverged in populations of progenitors such that *tbx2b* antagonizes *nrl* in developmental progenitors and *tbx2a* antagonizes *mafba* in Muller glia derived progenitors. It is interesting to speculate that the roles of *mafba* and *tbx2a* during regeneration are established during outgrowth from the CMZ. Therefore, these data indicate potential changes in the gene expression underlying photoreceptor specification dependent upon the origin of the progenitor, the neural epithelium, CMZ, or radial Muller glia. In chickens, where no *nrl* ortholog is maintained, a second Nrl family transcription factor, *MafA*, is thought to drive rod differentiation (Enright et el. 2015; Ochi et al. 2004; Kim et al. 2016) and *tbx2* is expressed in short cones which suggests a conserved function as seen in zebrafish (Enright et al. 2015). Other examples of subfunctionalization have been documented in the zebrafish retina primarily involving genes expressed in differentiated cells or genes involved in phototransduction (Renninger et al. 2011; Lerea et al. 1986; Haug et al. 2013; Lagman et al. 2015; Haug et al. 2024). It would be interesting to test if differential expression of the duplicate genes have led to adaptation of enzyme activity or binding partners better suited to their respective environments.

### *tbx2a* and *tbx2b* are redundant in other sensory systems

The shifting roles of *tbx2a* and *tbx2b* in retinal development suggests some level of selective pressure to retain both *tbx2a* and *tbx2b*. Homozygous *tbx2b^lor^*^/lor^ or *tbx2a^-/-^*are adult viable whereas *Tbx2* knockout in mice is embryonic lethal at embryonic day 8 likely due to cardiac defects (Harrelson et al. 2024). Analysis of cardiac phenotypes in *tbx2a* or *tbx2b* mutant zebrafish revealed subtle differences in embryonic development, opening the possibility of early stages of subfunctionalization (Sedletcaia and Evans, 2011; Tasnim et al. 2025). Single cell RNA sequencing data show co-expression of *tbx2a* and *tbx2b* during development of the auditory system and taste receptors (Farrell et al. 2018; Sur et al. 2023). Whereas no altered phenotypes were observed in single mutants, *tbx2a^-/-^; tbx2b^lor/lor^*larvae showed disrupted auditory and taste receptor development demonstrating genetic redundancy of *tbx2a* and *tbx2b*. The single otolith auditory phenotype observed in *tbx2a*; *tbx2b* double mutant zebrafish is consistent with cKO of *Tbx2* in mice, which resulted in disrupted inner hair cell differentiation (Kaiser et al. 2021; Kaiser et al. 2022; Garcia-Añoveros et al. 2022; Bi et al. 2024). We suspect the change in sws1 cone number does not significantly contribute to lethality (Bianco et al. 2011). Retinas of *tbx2b^lor/lor^* larvae contain very few sws1 cones randomly distributed across the retina. Double mutants did not display visual deficits and were able to inflate swim bladders. Unlike knockout of sws1 opsin which results in photoreceptor cell death or ablation of sws1 and lws cones which altered visual behaviors (Baden, 2024; Fornetto et al. 2025), we suspect retinal rewiring would accommodate for the absence of sws1 cones due to the cell fate changes (Saade et al. 2013; Yoshimatsu et al. 2014). The maintenance of *tbx2a* and *tbx2b* could protect against haploinsufficiency and cell fate changes from sws1 cone to rod to preserve sws1 cones necessary for prey capture.

## Conclusion

We propose a model whereby the expression of transcriptional repressors guides progression of multipotent progenitors through a hierarchy of competency states by antagonizing the activity of transcriptional activators, much like a series of locks and gates (Figure 7). This model is necessary because in zebrafish early multipotent progenitors co-express the transcriptional activators *crx*, *otx2*, *thrβ2* and *nr2e3*, amongst other, and possess the potential to adopt any one of a number of photoreceptor fates. Yet differentiation still occurs in an orderly, highly conserved sequence. We propose that repressors act as gates, initially preventing the multipotent progenitors from expressing multiple fates then allowing adoption of a unique cell fate or the progression to the next, lower-level competency state. In the multipotent progenitor, low level expression of *samd7* and *tbx2a/b*, and low-level notch signaling maintain the undifferentiated state. Derepression of *thrβ2* and *samd7* drives the adoption of an lws-cone fate while repressing later fates (Volkov et al. 2024). Notch signaling in neighboring cells represses *thrβ2* (Hedge et al. 2008). At the same time, decreased *samd7* expression derepresses *tbx2a/b* expression progressing the progenitor to the next state, or lock, as a bipotential progenitor with the ability to adopt an sws1 cone or rod fate (Volkov et al. 2024). *tbx2a/b,* serves as a gate to repress *nr2e3/nrl* function and the rod fate, while expression of a yet to be defined factor(s) specifies the sws1 cone fate. As *tbx2a/b* expression decreases in the remaining progenitors, *nr2e3/nrl* represses cone-specific genes and drives the rod fate (Mears et al. 2001; Oh et al. 2007; Oh et al. 2008; Oel et al. 2020). The rapid development and short delay between cell cycle exit and opsin expression necessitates the activity of factors that suppress the differentiation program in mitotic cells primed to differentiate into a specific cell fate.

**Figure 7:**
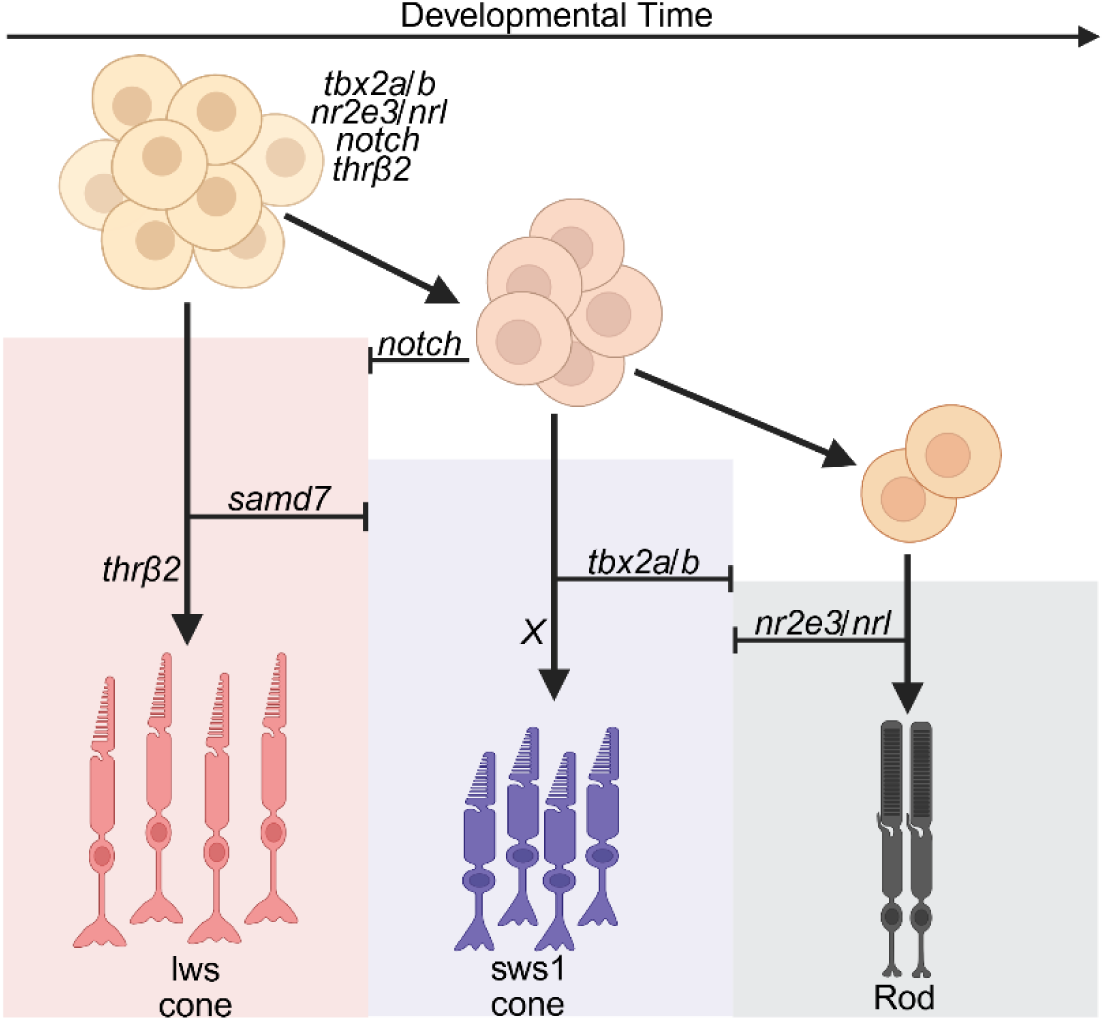
Model for the photoreceptor fate specification. A common progenitor expresses multiple transcriptional activators. *thrβ2* drives the lws cone fate. *samd7* represses *tbx2a/b* preventing the progression into the next competency state. Notch inhibits *thrβ2*, advancing the progenitor to a lower competency level. *tbx2a/b* represses the rod fate, while a yet to be defined factor drives the sws1 cone fate. *nr2e3* represses cone specific genes and drives the expression of *nrl* driving the remaining photoreceptor progenitors to adopt a rod fate.

## Acknowledgements

We thank Tyson von Scherrer, Kealyn Bowie, and Beija Gore for fish husbandry, the Florida State University DNA Sequencing Laboratory, Molecular Cloning Laboratory and Biological Imaging facility. We thank undergraduates Lucas Cruz, William Clement, Kennady Lange, and Christian Noble for prepping samples for genotyping. This work was supported by National Institutes of Health Grant R01EY030633.

## Methods

### Fish Husbandry

Zebrafish (Danio rerio) wildtype AB and TL strains were used throughout all experiments. *Tg(−5.5sws1:EGFP)* were used to fluorescently tag sws1 cones (Takechi et al. 2003). The *tbx2b^p25bbtl^ (lots-of-rods, tbx2b^lor^)* mutant was previously isolated and characterized (Alvarez-Delfin et al. 2009, Saade et al. 2013). All procedures were approved by the Florida State University (FSU) Institutional Animal Care and Use Committee, AUCU Protocol #PROTO202000004. Larval and adult animals were anesthetized using Syncaine and euthanized in ice water.

### CRISPR/Cas9 Mutagenesis and Transgenic Generation

All guides were designed using IDT. Guides were selected based upon low off-target scores and location within the exons. Guides were precomplexed with Cas9 (IDT) at a concentration of 5 μM. After precomplexing, guides for the same target gene were mixed at a 1:1 ratio (2 guides for *tbx2a*, 2 guides for *thrβ2*, 3 guides for *nrl*, 1 guide for *six7,* Table S1). Single cell stage embryos were injected with 3-5 nL of ribonucleoprotein (RNP) solution. Embryos were raised and inbred to produce F_1_s. F_1_s were screened for mutation using PCR amplification with primers located up and downstream of the guide landing zone (Table S2). PCR samples were purified and sent for sequencing at the FSU Biological Science Core. Mutant zebrafish were crossed based upon predicted protein sequence disruption. F_2_s were fixed and screened for changes in photoreceptor type and number.

An *EGFP:tbx2b* fusion protein was overexpressed in cones using the *alpha transducin* (*gnat2*) promoter (Kennedy et al. 2007, Kulisz and Simon, 2008, Charest et al. 2020). Single-cell stage wildtype and *tbx2* double mutant embryos were injected with transposase mRNA and a plasmid containing a (*Tol2-gnat2:EGFP-tbx2b-polyA*) cassette and a *cmlc2:EGFP* cassette that drive heart specific expression of EGFP (Kwan et al. 2007). Embryos were screened for EGFP positive hearts to confirm transgenesis. Immunohistochemistry was performed at 5 days post fertilization (dpf) to determine the fate of EGFP expressing cone cells.

### Morpholino Knockdown

Morpholino (MO) knockdown of thrb2 was done following Suzuki et al. Briefly, a MO specific to the retinal specific splice site (5’-TCTAGAACTTGCAATACCTTTCTTA-3’) of thrb2 was injected into one-cell stage wildtype, *from beyond (tbx2b^fby^)*, and *tbx2b^lor^* embryos (Genetools) (2013). A standard control MO was used as a negative control.

### Quantitative PCR

qPCR was performed following procedures from Sontolongo-Lopez et al. with an additional purification step (2016). Briefly, RNA from whole 6 dpf larvae pooled into groups of 30 was extracted using TRIzol (Invitrogen). Tissue was homogenized and chloroform was added for phase separation. The aqueous phase was washed with alcohol. RNA was purified using an RNA purification kit (Xymo) and following manufacturer’s protocol with in-column DNase treatment. Primers for qPCR are listed in Supplementary Table 3. The FSU Biological Core performed qPCR using a SYBR Green qPCR Master Mix kit and protocols (ThermoFisher). βactin was used to normalize for fold expression changes and relative quantity was calculated following standard procedure (Livak et al. 2001).

### Luciferase Reporter Assay Plasmids

2.3 kb containing tbx2b ORF was PCR-amplified and cloned into pCR2.1-TOPO, and the KpnI-XhoI fragment was subcloned into of the pCDNA3.1+ expression vector. Human Crx (h-Crx) and Xenopus laevis Nrl (xl-Nrl) cloned in pCS2+ vector, and the Xenopus rhodopsin promoter constructs were provided by Dr. Barry Knox (Mani et al. 2001; Whitaker and Knox, 2004; McIlvain and Knox, 2007). Rhod5500 contained the rhodopsin promoter fragment −5500/+41, cloned in the pGL3 basic vector (Promega, Madison WI). To generate the vector CMV-Luc, a MluI-HindIII fragment containing the CMV promoter was subcloned from pCDNA3.1+ into the pGL3 basic vector (Promega, Madison WI).

### Cell culture

Human embryonic kidney cells (HEK293T) were cultured in HyClone DMEM/High glucose medium (Gibco-Invitrogen, Life Technologies, Carlsbad CA), supplemented with 10% fetal bovine serum, 2 mM L-glutamine and antibiotics and kept at 37°C and 6% CO_2_ (Liang et al. 2011). Cells were transfected using OptiMEM reduced serum medium (Gibco-Invitrogen, Life Technologies, Carlsbad CA).

### Luciferase reporter assay

Luciferase assays were performed as previously described (Zhu et al. 2002). Briefly, subconfluent HEK293T cells grown in 24-well plates were transfected with DNA vectors using Lipofectamine 2000 according to manufacturer protocol (Invitrogen, Carlsbad CA). The amounts of plasmids used were: 100 ng of each promoter construct, increasing amounts of tbx2b or tbx2-RERE (0-100 ng), 100 ng of each of the activators xl-Nrl and/or h-Crx, and 10 ng of the Renilla luciferase pRL-TK vector as internal control. The correspondent empty plasmid was added to samples to equalize the amount of DNA in each transfection reaction. Luciferase activity was detected 24 hours after transfection using the Dual-Luciferase Reporter Assay System (Promega, Madison WI) according to manufacturer protocol. Relative luciferase activity is the ratio between the Firefly luciferase and the Renilla luciferase values. Results were presented as fold change relative to the samples without activation.

### Imaging and Image Analysis (Immunohistochemistry)

Whole-mount 5 and 6-dpf embryo and frozen sections (8-10 microns) were labelled as previously described (Morris et al. 2008; Saade et al. 2013). Fluorescence microscopy was performed on a Nikon Spinning Disk confocal equipped with either a 20X water emersion objective (NA 0.95) or 40X water emersion objective (NA 1.15) and super-resolution. The following mouse monoclonal antibodies were used: 4C12 antibody (1:150) was used as a rod marker. Arrestin 3A (zpr1) antibody (1:100, ZIRC) was used to label red and green cone cell bodies (Renninger et al. 2011). Zpr3 (1:100, Zirc) was used to label rhodopsin and rhodopsin-2 (Hu et al. 2024). Commercially available rabbit polyclonal antibodies sws2 opsin (1:500, Kerafast) and sws1 opsin (1:120, LS Biology) were used to label short-wavelength cone outer segments. Chicken anti-GFP (1:500, Aves Lab, Cat: GFP-1020) was used to enhance GFP label. Click-It EdU Alexa Fluor 546 Imaging kit (Invitrogen) was used to label proliferating cells with EdU incorporated into DNA during S-phase (Salic et al. 2008). All figures were generated using https://BioRender.com. Images were rotated to maintain a dorsal ventral orientation. All images had the brightness adjusted to 200% for visualization of antibody labeling.

### Light Damage and EdU Injections

Age and size matched male and female adult wildtype and mutant zebrafish with the −5.5kbsws1:EGFP transgenes or both the −5.5kbsws1:EGFP and *Tg(pou4f3:GAP-GFP)* were dark adapted for 7 days. Following dark adaptation, fish are placed in 120,000 lux light for 30 minutes, then moved into 20,000-60,000 lux constant white light for 96 hours (Thomas and Thummel, 2013). 10 mM EdU (Invitrogen) was injected intraperitoneal between the pelvic and anal fins at 36, 60, and 84 hours post light exposure to label proliferating Muller glia derived progenitors (Campbell et al. 2021). A subset (n=5) of each genotype were dissected after 96 hours of light damage to confirm photoreceptor cell death. A second group was allowed to recover for 10 days to observe regeneration of photoreceptors. 10-micron thick frozen sections were immunolabelled using standard procedures. Photoreceptor specific antibodies and Click-It EdU labelling was performed to label newly regenerated photoreceptors. A two-way ANOVA with a Bonferroni’s post-hoc was performed for statistical analysis.

### Quantitative Analysis

Eyes were removed from immunolabelled 5 dpf wholemount larva and mounted on a coverslip lens down in 0.8% LMP agarose in 50% glycerol. A 3500 μm^2^ region opposite the ventral patch dorsal to the optic nerve head was counted using FIJI Cell Counter to quantify photoreceptor number. 8-10 μm sections of larva and adult sections were quantified in centrally located sections. 60-day old retinas for wildtype (n=5), *tbx2a^fl18/fl20^* (n=5*), tbx2a^fl18/fl20^*; *tbx2b^lor/+^* (n=5), and *tbx2b^lor/lor^* (n=5) were imaged and loaded into FIJI software. The outer nuclear layer was hand traced using the segmented line feature. The fit spline feature was used to cover the entire width of the outer nuclear layer and outer segments of the sws1 cones. Images were then straightened with the dorsal retina located to the left and the ventral retina located on the right. Images were then thresholded and exported. Python (version 3.12) was then used to scan a sliding 30 micron wide by 50 high window with a 50% overlapping area the length of the outer nuclear layer. Images were then normalized to the brightest point of each individual fish. Average fluorescent intensity was plotted for each genotype and the average of all genotypes.

Statistical analysis was performed using GraphPad Prism V.11.0. One-way ANOVA with a Bonferroni’s post-hoc was performed on larval datasets comparing differences across genotypes. Unpaired Student’s t-test were performed on CRISPant datasets. Two-way ANOVA with a Bonferroni’s post-hoc was performed on the light damage dataset comparing across treatment (LD) and genotypes. A p-value <0.05 was considered statistically significant. One-tail Mann Whitney test was used to analyze changes in cone opsin expression in qPCR. A two-tail Mann Whitney test was used to analyze changes in Rhodopsin expression, as we did not expect to see changes in expression due to cone specific promoter.

## Supplementary Figures and Tables

**Supplementary Figure 1:**
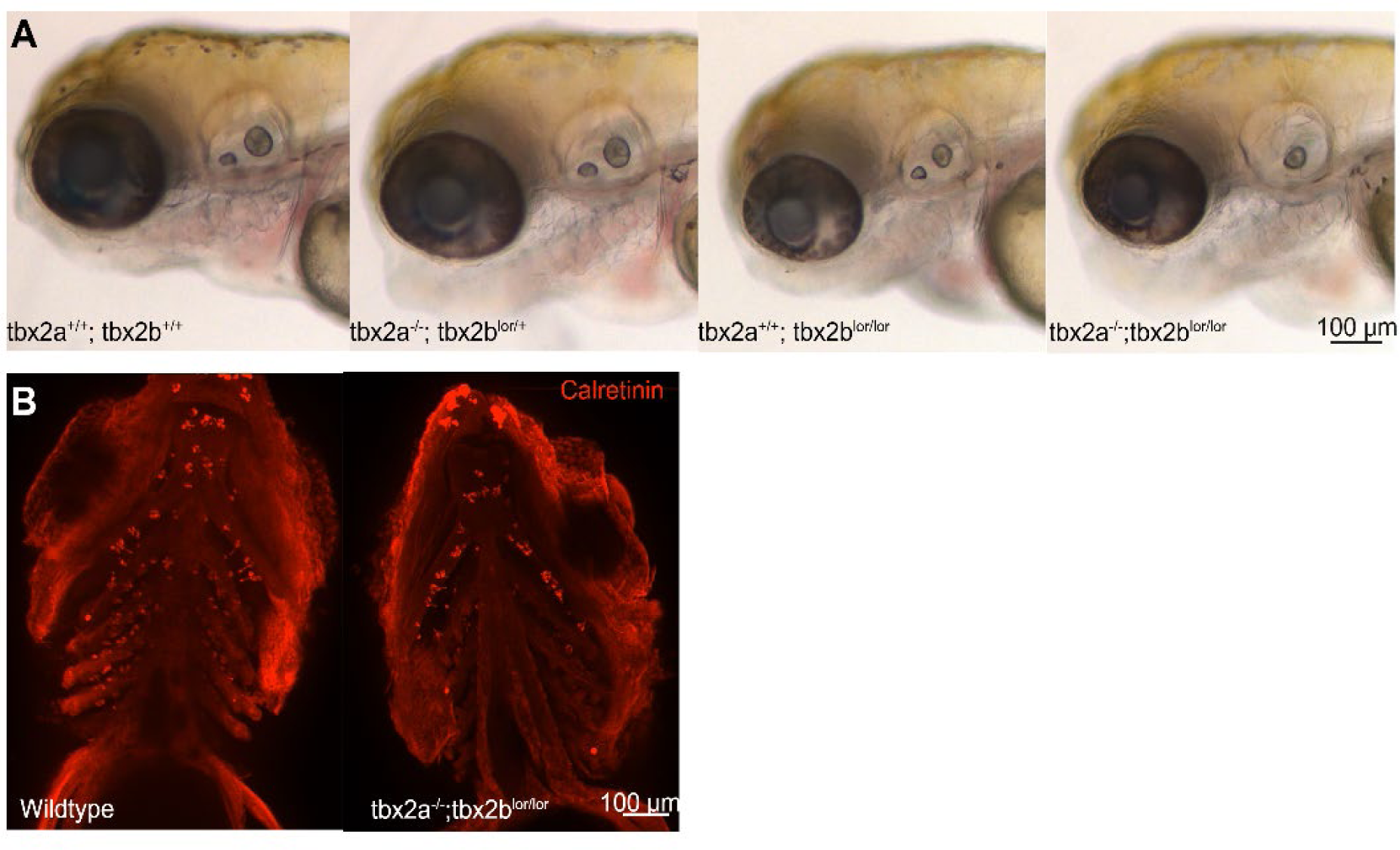
*tbx2a^-/-^; tbx2b^lor/lor^* larvae display auditory (A) and Calretinin+ cell distribution phenotypes(B).

**Supplementary Figure 2:**
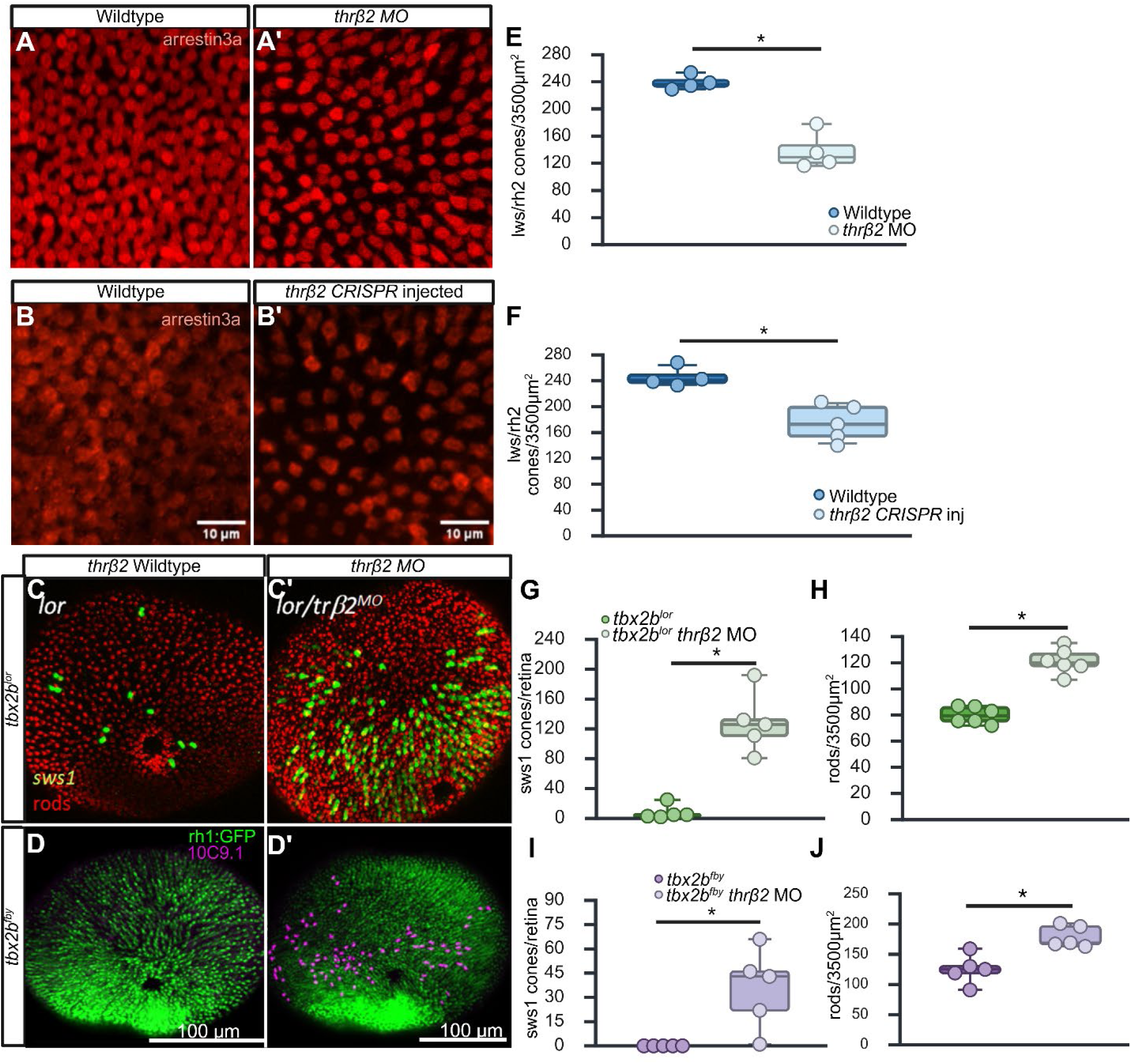
*thrβ2* targeting using a morpholino or CRISPR/Cas9 leads to a significant reduction in rh2/lws cones and results in a significant increase in both sws1 cones and rods in *tbx2b* mutants. A-B) Wholemount immunolabeling for arrestin3a in 5 dpf wildtype embryos. Distribution of rh2/lws cones just dorsal the optic nerve in 5 dpf larvae. A’-B’) Targeting of thrb2 using a morpholino or CRISPR/Cas9 resulted in a significant reduction in rh2/lws cones, attributed to the loss of lws cones. C) *tbx2b^lor/lor^* and D) *tbx2b^fby/fby^* animals have a reduced number of sws1 cones and a concomitant increase in rods. C’) tbx2b^lor/lor^ and D’) *tbx2b^fby/fby^* animals injected with a morpholino specific to thrb2 resulted in a significant increase in both sws1 cones (purple) and rods (green). E-F) Quantification of lws/rh2 cones in morpholino (E): wildtype (n=4, 239+/-10.5), *thrβ2* MO (n=138+/-27.8) and *thrβ2* CRISPR (F) injected: wildtype (n=4, 245+/-15.5), thrb2 inj. (n=5, 175+/-28.5). G-J) Quantification of sws1 cones per retina (G: tbx2b^lor/lor^ (n=5, 8+/-9.6), tbx2b^lor/lor^ thrb2 MO(n=5, 128+/-40.6))+ I: tbx2b^fby/fby^ (n=5, 0), tbx2b^fby/fby^ thrb2 MO (n=5, 36+/-24.8) and rods/3500 µm^2^ (H: tbx2b^lor/lor^ (n=6, 80+/-6.4), tbx2b^lor/lor^ thrb2 MO (n=6, 121+/-9.6)+J: tbx2b^fby/fby^ (n=5, 125+/-24.3), *tbx2b^fby/fby^ thrb2* MO (n=5, 179+/-17.8) in morpholino injected animals. Student’s unpaired t-test p<0.05.

**Supplementary Figure 3:**
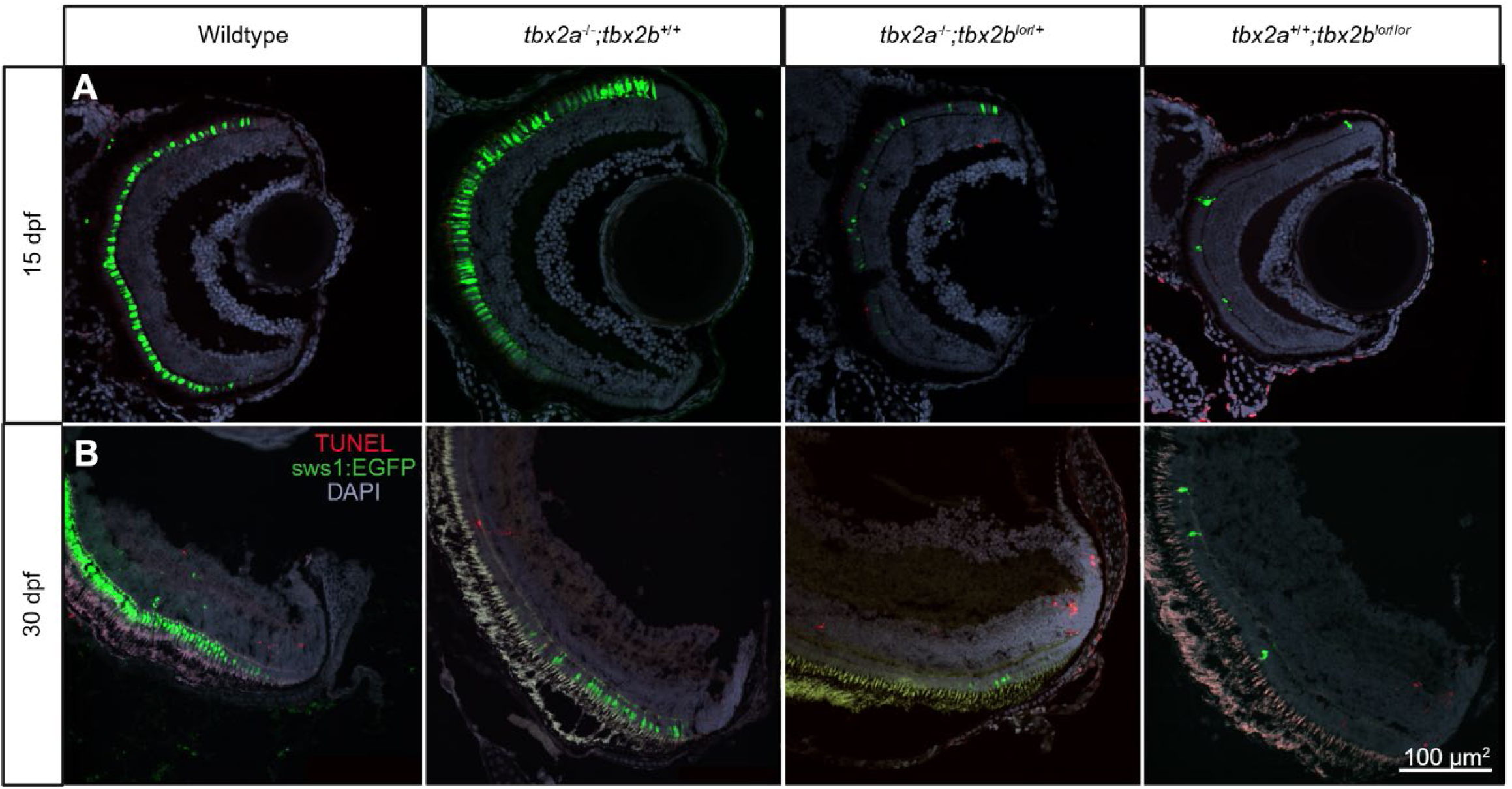
Cell death was not observed at the boundary of sws1 cones. A-B) Immunolabeling for sws1EGFP (green) and TUNEL (Red) at 15 dpf (A) and 30 dpf (B) across genotypes. No difference was observed in the amount of TUNEL+ cells across genotypes

**Supplementary Table 1:**
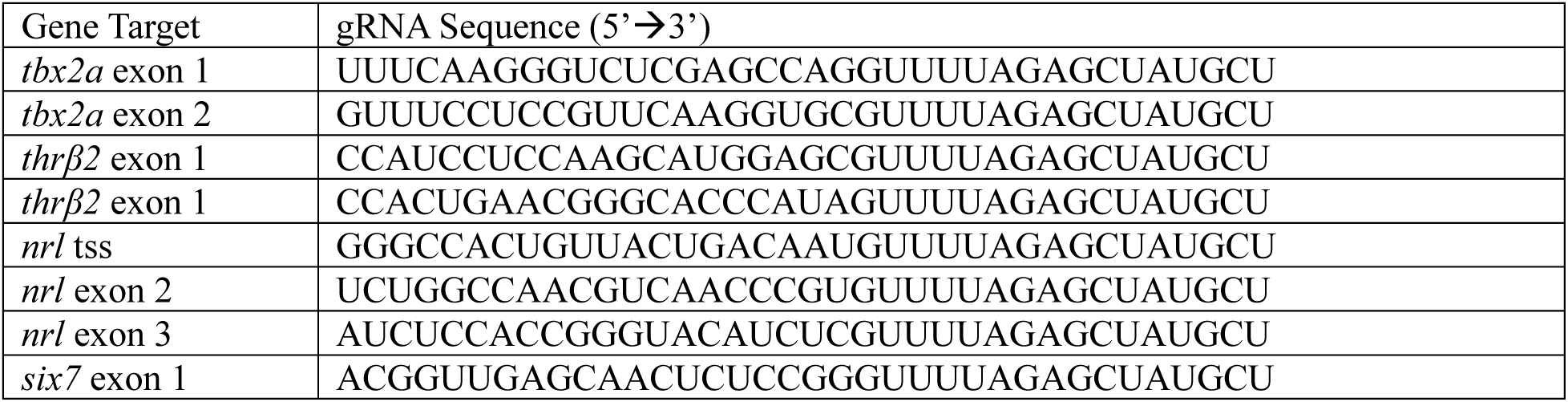
gRNA sequences used for CRISPR/Cas9 targeting.

**Supplementary Table 2:**
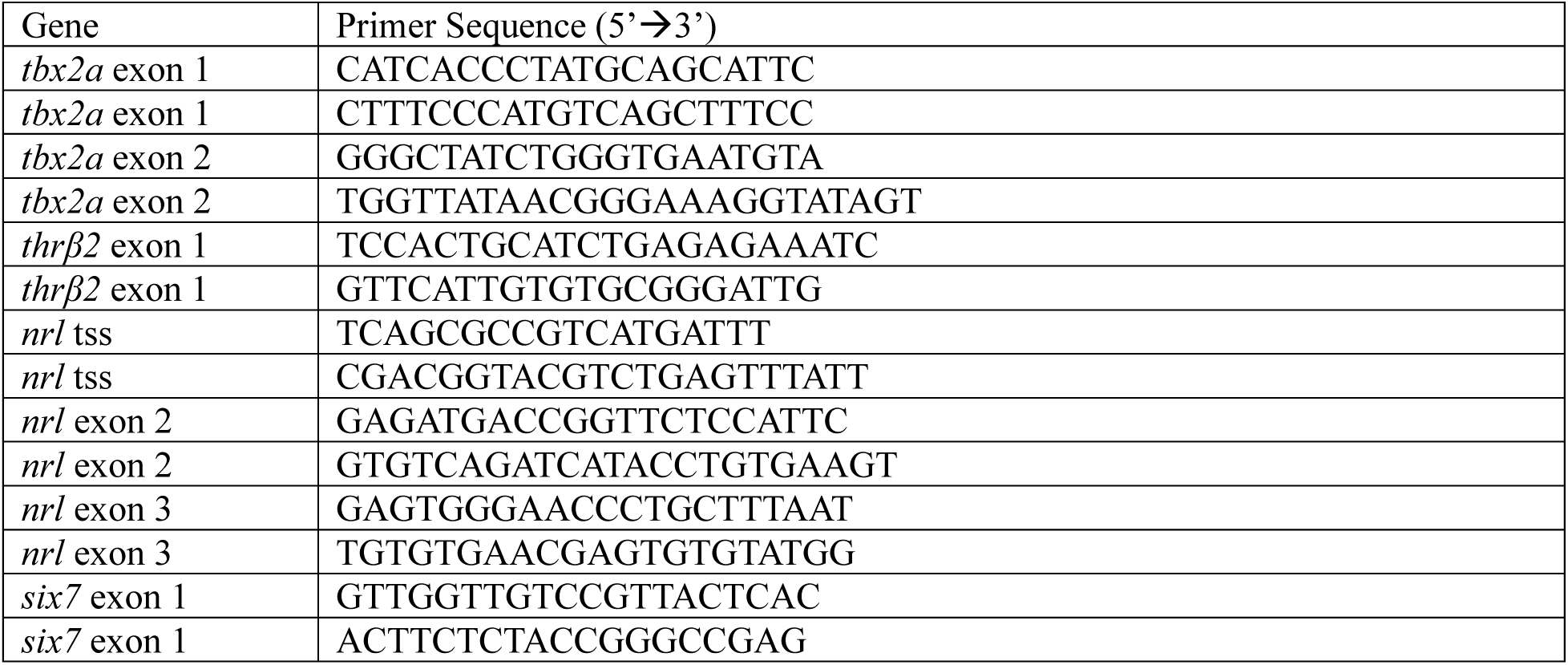
DNA primer sequences used for screening lesions across different loci.

**Supplementary Table 3:**
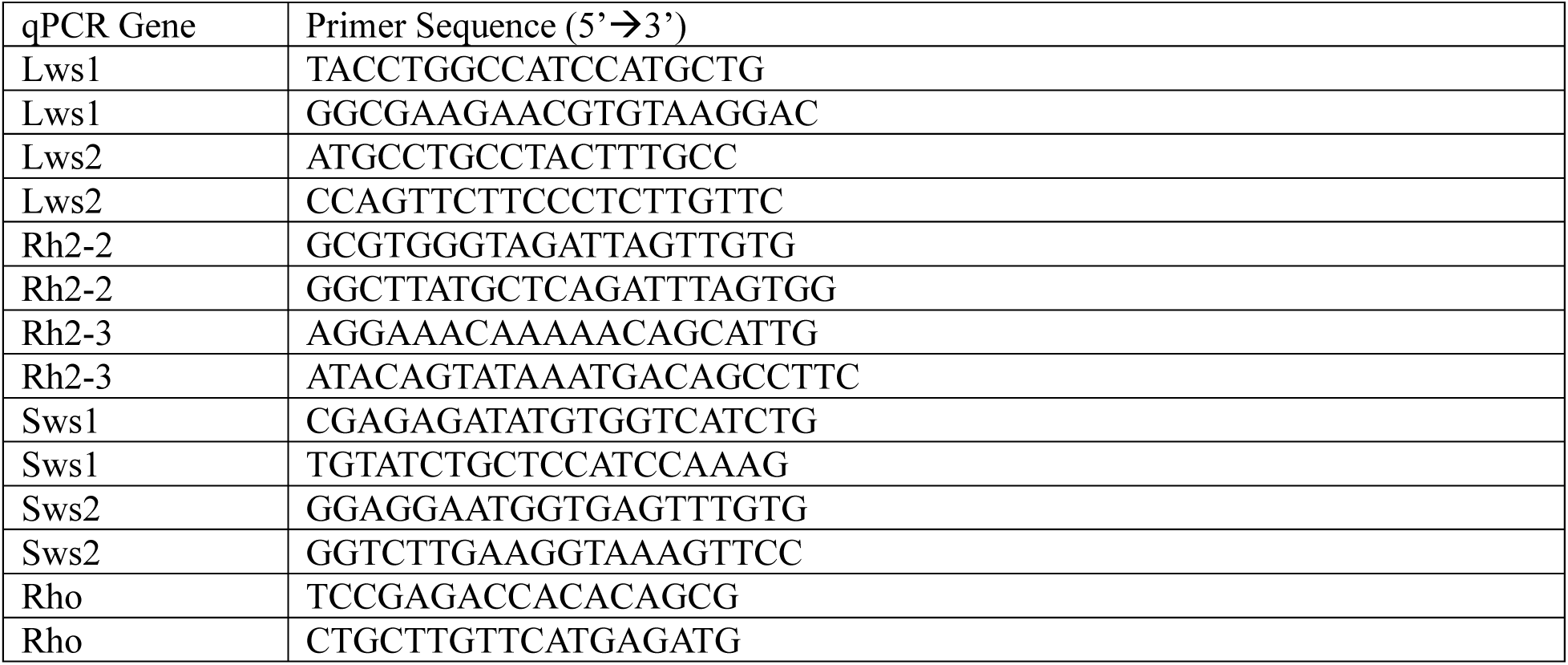
Primer sequences used for qPCR.

