## Supplementary Table 3 for "Subfunctionalization of *tbx2* paralogues during photoreceptor cell specification in zebrafish"

| qPCR Gene | Primer Sequence (5'→3') |
| --- | --- |
| Lws1 | TACCTGGCCATCCATGCTG |
| Lws1 | GGCGAAGAACGTGTAAGGAC |
| Lws2 | ATGCCTGCCTACTTTGCC |
| Lws2 | CCAGTTCTTCCCTCTTGTTC |
| Rh2-2 | GCGTGGGTAGATTAGTTGTG |
| Rh2-2 | GGCTTATGCTCAGATTTAGTGG |
| Rh2-3 | AGGAAACAAAAACAGCATTG |
| Rh2-3 | ATACAGTATAAATGACAGCCTTC |
| Sws1 | CGAGAGATATGTGGTCATCTG |
| Sws1 | TGTATCTGCTCCATCCAAAG |
| Sws2 | GGAGGAATGGTGAGTTTGTG |
| Sws2 | GGTCTTGAAGGTAAAGTTCC |
| Rho | TCCGAGACCACACAGCG |
| Rho | CTGCTTGTTCATGAGATG |

Supplementary Table 3: Primer sequences used for qPCR.
