## Supplementary Table 2 for "Subfunctionalization of *tbx2* paralogues during photoreceptor cell specification in zebrafish"

| Gene | Primer Sequence (5'→3') |
|---|---|
| *tbx2a* exon 1 | CATCACCCTATGCAGCATTC |
| *tbx2a* exon 1 | CTTTCCCATGTCAGCTTTCC |
| *tbx2a* exon 2 | GGGCTATCTGGGTGAATGTA |
| *tbx2a* exon 2 | TGGTTATAACGGGAAAGGTATAGT |
| *thrβ2* exon 1 | TCCACTGCATCTGAGAGAAATC |
| *thrβ2* exon 1 | GTTCATTGTGTGCGGGATTG |
| *nrl* tss | TCAGCGCCGTCATGATTT |
| *nrl* tss | CGACGGTACGTCTGAGTTTATT |
| *nrl* exon 2 | GAGATGACCGGTTCTCCATTC |
| *nrl* exon 2 | GTGTCAGATCATACCTGTGAAGT |
| *nrl* exon 3 | GAGTGGGAACCCTGCTTTAAT |
| *nrl* exon 3 | TGTGTGAACGAGTGTGTATGG |
| *six7* exon 1 | GTTGGTTGTCCGTTACTCAC |
| *six7* exon 1 | ACTTCTCTACCGGGCCGAG |

Supplementary Table 2: DNA primer sequences used for screening lesions across different loci.
