## Supplementary Table 1 for "Subfunctionalization of *tbx2* paralogues during photoreceptor cell specification in zebrafish"

| Gene Target | gRNA Sequence (5'→3') |
| --- | --- |
| *tbx2a* exon 1 | UUUCAAGGGUCUCGAGCCAGGUUUUAGAGCUAUGCU |
| *tbx2a* exon 2 | GUUUCCUCCGUUCAAGGUGCGUUUUAGAGCUAUGCU |
| *thrβ2* exon 1 | CCAUCCUCCAAGCAUGGAGCGUUUUAGAGCUAUGCU |
| *thrβ2* exon 1 | CCACUGAACGGGCACCCAUAGUUUUAGAGCUAUGCU |
| *nrl* tss | GGGCCACUGUUACUGACAAUGUUUUAGAGCUAUGCU |
| *nrl* exon 2 | UCUGGCCAACGUCAACCCGUGUUUUAGAGCUAUGCU |
| *nrl* exon 3 | AUCUCCACCGGGUACAUCUCGUUUUAGAGCUAUGCU |
| *six7* exon 1 | ACGGUUGAGCAACUCUCCGGGUUUUAGAGCUAUGCU |

Supplementary Table 1: gRNA sequences used for CRISPR/Cas9 targeting.
