## Supplementary Figure 3 for "Subfunctionalization of *tbx2* paralogues during photoreceptor cell specification in zebrafish"

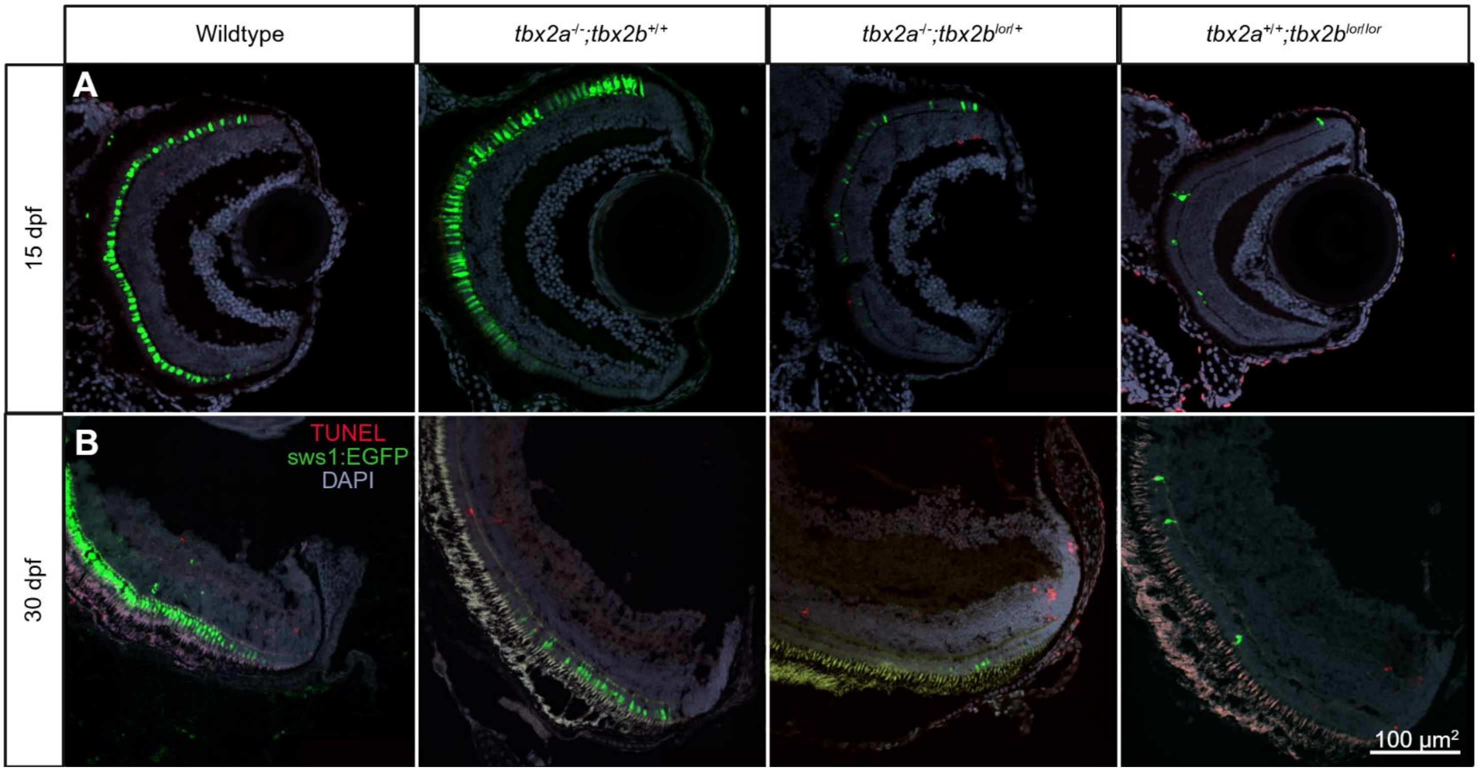


Supplementary Figure 3: Cell death was not observed at the boundary of sws1 cones. A-B) Immunolabeling for sws1EGFP (green) and TUNEL (Red) at 15 dpf (A) and 30 dpf (B) across genotypes. No difference was observed in the amount of TUNEL+ cells across genotypes.
