## Supplementary Figure 2 for "Subfunctionalization of *tbx2* paralogues during photoreceptor cell specification in zebrafish"

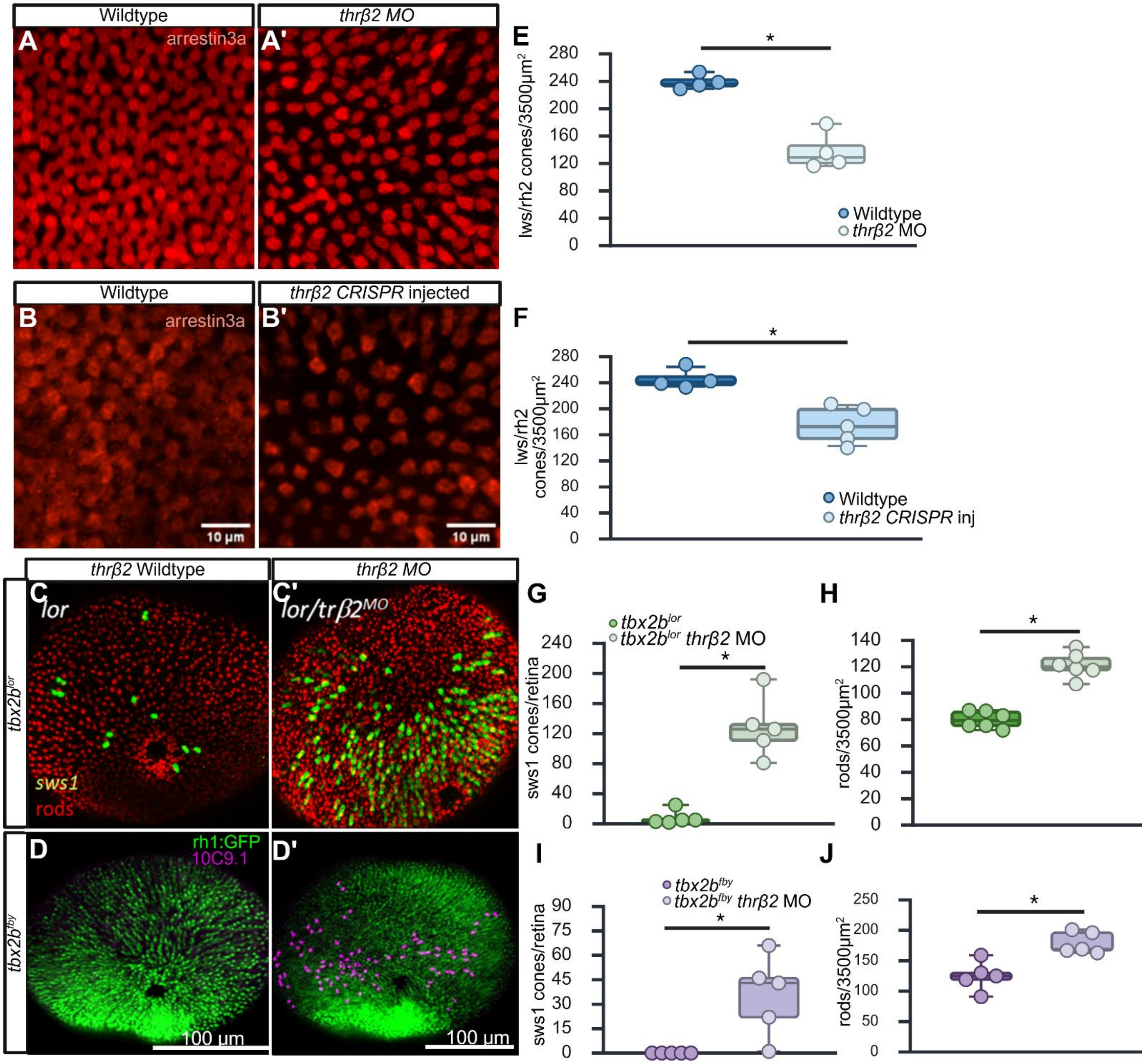

Supplementary Figure 2: *thrβ2* targeting using a morpholino or CRISPR/Cas9 leads to a significant reduction in rh2/lws cones and results in a significant increase in both sws1 cones and rods in *tbx2b* mutants. A-B) Wholemount immunolabeling for arrestin3a in 5 dpf wildtype embryos. Distribution of rh2/lws cones just dorsal the optic nerve in 5 dpf larvae. A'-B') Targeting of thrb2 using a morpholino or CRISPR/Cas9 resulted in a significant reduction in rh2/lws cones, attributed to the loss of lws cones. C) *tbx2b*$^{lor/lor}$ and D) *tbx2b*$^{fby/fby}$ animals have a reduced number of sws1 cones and a concomitant increase in rods. C') tbx2b$^{lor/lor}$ and D') *tbx2b*$^{fby/fby}$ animals injected with a morpholino specific to thrb2 resulted in a significant increase in both sws1 cones (purple) and rods (green). E-F) Quantification of lws/rh2 cones in morpholino (E): wildtype (n=4, 239+/-10.5), *thrβ2* MO (n=138+/-27.8) and *thrβ2* CRISPR (F) injected: wildtype (n=4, 245+/-15.5), thrb2 inj. (n=5, 175+/-28.5). G-J) Quantification of sws1 cones per retina (G: tbx2b$^{lor/lor}$ (n=5, 8+/- 9.6), tbx2b$^{lor/lor}$ thrb2 MO(n=5, 128+/-40.6))+ I: tbx2b$^{fby/fby}$ (n=5, 0), tbx2b$^{fby/fby}$ thrb2 MO (n=5, 36+/-24.8) and rods/3500 μm$^2$ (H: tbx2b$^{lor/lor}$ (n=6, 80+/-6.4), tbx2b$^{lor/lor}$ thrb2 MO (n=6, 121+/-9.6)+J: tbx2b$^{fby/fby}$ (n=5, 125+/-24.3), *tbx2b*$^{fby/fby}$ *thrb2* MO (n=5, 179+/-17.8) in morpholino injected animals. Student's unpaired t-test $p<0.05$.
