## Supplementary Figure 1 for "Subfunctionalization of *tbx2* paralogues during photoreceptor cell specification in zebrafish"

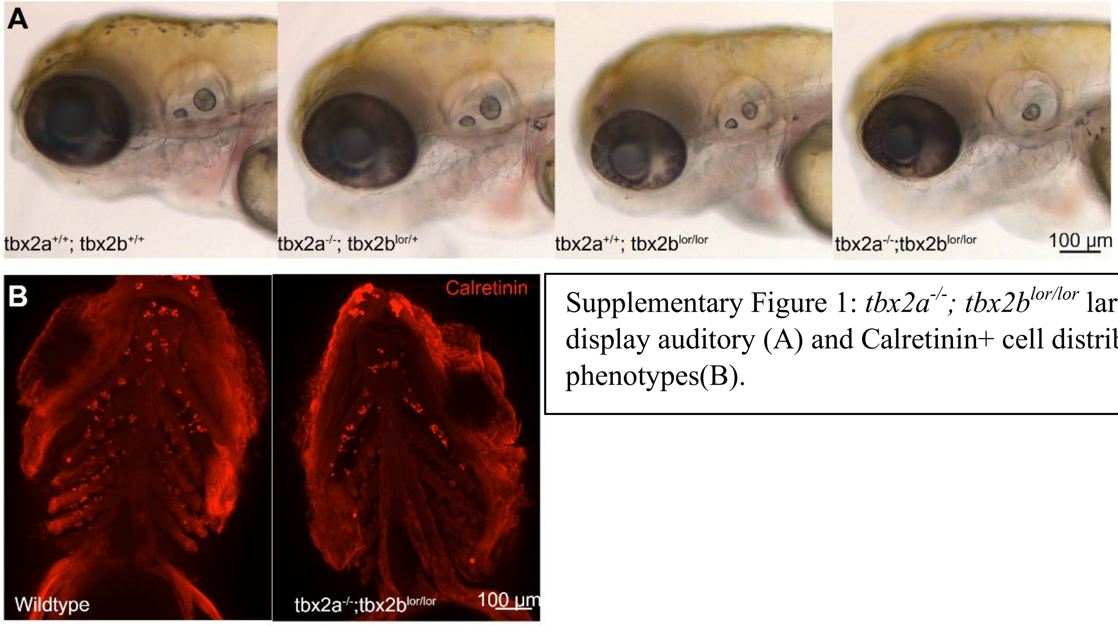


Supplementary Figure 1: *tbx2a*$^{-/-}$*; tbx2b*$^{lor/lor}$ larvae display auditory (A) and Calretinin+ cell distribution phenotypes(B).
